# Dogs, but not wolves, lose their sensitivity towards novelty with age

**DOI:** 10.1101/405480

**Authors:** Christina Hansen Wheat, Wouter van der Bijl, Hans Temrin

**Author notes:** Correspondence: Christina Hansen Wheat.

## Abstract

Selection on behavioural traits holds a prominent role in the domestication of animals, with reductions in fear behaviour considered to be a key component. Specifically, there is a general assumption that domesticated species express reduced fear and reactivity towards novel stimuli compared to their ancestral species. However, very few studies have explicitly tested this proposed link between domestication and reduced fear responses. Of the limited number of studies experimentally addressing the alterations of fear during domestication, the majority have been done on canids. Previous work in foxes, wolves and dogs has led to the suggestion that decreased expression of fear in domesticated animals is linked to a domestication driven delay in the first onset of fearful behaviour during early ontogeny. Thus, wolves are expected to express exaggerated fearfulness earlier during ontogeny compared to dogs. However, while adult dogs are less fearful towards novelty than adult wolves and wolf-dog hybrids, consensus is lacking on when differences in fear expression arise in wolves and dogs. Here we present the first extended examination of fear development in hand-raised dogs and European grey wolves, using repeated novel object tests from six to 26 weeks of age. Contrary to expectations, we found no evidence in support of an increase in fearfulness in wolves with age or a delayed onset of fear response in dogs compared to wolves. Instead, we found that dogs strongly reduced their fear response in the period between six and 26 weeks of age, resulting in a significant species difference in fear expression towards novelty at 26 weeks. Critically, as wolves did not differ in their fear response towards novelty over time, the detected species difference was caused solely by a progressive reduced fear response in dogs. Our results thereby suggest that species differences in fear of novelty between wolves and dogs are not caused by a domestication driven shift in the first onset of fear response. Instead, we suggest that a loss of sensitivity towards novelty with age in dogs causes the difference in fear expression towards novelty in wolves and dogs.

## Introduction

Humans have successfully domesticated a wide range of plants and animals and abundant evidence demonstrates how domesticated species express dramatically altered phenotypes compared to their wild counterparts (Driscoll et al. 2009). In animals, selection on behavioural traits had a prominent role in creating the domesticated phenotype (Belyaev et al. 1985; Trut 1999). In wild animal populations, fear is a key behaviour as a timely and proper response to novelty (e.g. flight response versus exploration) has direct fitness consequences (Boissy 1995; Weidenmayer 2009). In contrast, in domesticated animals living in human-controlled environments, strong fear responses and high reactivity towards novelty are undesirable traits (Leiner and Fendt 2011), and selection for docility (i.e. tameness) and against fearfulness was likely a key component in the successful domestication of animals (Belyaev et al. 1985; Trut 1999). Consequently, there is now a general assumption that domesticated species express reduced flight distances and reactivity towards novel stimuli (Zeder 2012) compared to their ancestral species. However, though good evidence exists that cortisol secretion and brain structures associated with fear responses have been significantly reduced in domesticated animals (Kruska 1988; Trut et al. 2009), excessive fear behaviour prevails in various domesticated species (Hemsworth et al. 1996), including rabbits (Csatádi et al. 2005), chickens (Jones and Waddington 1992), dogs (Döring et al. 2009) and horses (Christensen et al. 2008). These discrepancies impair our understanding of how the expression of fear has changed during animal domestication and this shortcoming is further complicated by the fact that very few studies have explicitly tested the proposed link between domestication and reduced fear responses.

In wild populations appropriate fear responses are formed and modified throughout ontogeny, during which juvenile animals gradually combine individual experience and social information, thereby developing the ability to discriminate between threatening and neutral stimuli (Scott and Fuller 1965; Griffin 2004; Weidenmayer 2009). Ontogeny has been modified in several ways during domestication and compared to ancestral species, domesticated animals express altered developmental rates, a phenomenon known as heterochrony (Goodwin et al. 1997; Price 1999; Dobney and Larson 2006). Specifically, domesticated animals express accelerated and/or delayed onsets of various ontogenetic stages, such as earlier sexual maturation and the retention of juvenile traits into adulthood (Morey 1994; Price 1999; Coppinger et al. 1987; Crockford 2002). Heterochrony has been suggested to affect behavioural ontogeny by prolonging the sensitive period (Martin 1978; Belyaev et al. 1985; Gariépy et al. 2001; Wilkins et al. 2014), an important period during behavioural development in which the juvenile animal is particularly sensitive to imprint on and form social bonds with conspecifics (Scott 1962; Scott and Fuller 1965; Freedman et al. 1961; Coppinger and Coppinger 2001). During the sensitive period juvenile animals show increased exploratory behaviour, as they readily approach novel stimuli and thereby learn about and socialize with their environment (Morrow et al. 2015). Importantly, the end of the sensitive period is marked by a progressive increase in fear and decreased exploration of novelty (Freedman et al. 1961; Belyaev et al. 1985). Based primarily on the findings in a long-term selection study on silver foxes (*Vulpes vulpes*), it has been suggested that domestication causes a shift in the sensitive period resulting in a delayed onset of fearful response in domesticated compared to non-domesticated animals (Belyaev et al. 1985; Trut, et al. 2004) (but see also Coppinger and Coppinger, 2001). While this might indicate that differences in fear expression between domesticated and non-domesticated animals arise already during early ontogeny, only a very limited body of studies have experimentally compared the ontogeny of fear in wild and domestic species under controlled conditions and with ambiguous results (Bilkó and Altbäcker 2000; Lord 2013). Therefore, it remains largely an open question whether the ontogeny of fear and the sensitive period have been altered by domestication.

The domestic dog (*Canis familiaris*) is an excellent study species when addressing questions about how domestication has affected behavioural ontogeny. Domestication of the dog from the grey wolf (*Canis lupus*) occurred at least 15,000 years ago (Driscoll et al. 2009), making the dog the first domesticated species and with the ancestral species extant, the opportunities for comparisons are ideal (Price 2002). Studies of behavioural ontogeny in dogs have largely focused on the sensitive period, and fear of novelty in the dog puppy has traditionally been reported to manifest at eight weeks of age and continually increase onward (Scott and Marston 1950; Scott 1958; Freedman et al. 1961; Scott and Fuller 1965). However, recent evidence suggests that the development of fear might be highly breed-specific and subject to considerable variation (Morrow et al. 2015), thereby highlighting substantial gaps in our knowledge of the ontogeny of fear in dogs. In wolves, consensus on when fear behaviour is established is lacking, with the onset of fearful response reported to occur as varied as four to eight weeks of age across studies (Scott and Marston 1950; Fentress 1967; Wooply and Ginsburg 1967; Fox 1972; Zimen 1987; Lord 2013). The ambiguity of these wolf studies is further complicated by the fact that the majority of them were conducted over a short period of time and/or focused on isolated individuals or single litters, thereby limiting our ability to generalize from these findings. Additionally, a recent study found no difference in fear related behaviours or the latency to make contact with a novel object in six and eight week old wolves and dogs(Marshall-Pescini et al. 2017), thereby suggesting that wolves might not express fear towards novelty at an earlier age than dogs. Thus, while adult wolves (Moretti et al. 2015) and wolf-dog hybrids (Hansen Wheat et al. 2018) are more fearful of novelty than dogs, the question of when during development species differences in fear expression is established remains unresolved. Furthermore, both juvenile and adult wolves explore and interact with novel objects more than similar aged dogs (Moretti et al. 2015; Marshall-Pescini et al. 2017), and adult dogs have been reported to be less likely to approach a novel object than wolves (Moretti et al. 2015). While these findings have been interpreted as less interest in novelty, and not fear, in dogs compared to wolves (Moretti et al. 2015), more studies are needed to tease these components apart and provide more detailed insight into how, and at which developmental stage, domestication changes fear expression in wolves and dogs.

The lack of consensus across studies comparing wolves and dogs to uncover implications of domestication illustrates a fundamental challenge in this field, namely the combination of limited animal availability and the enormous effort necessary to hand-raise, socialize and test acquired animals. These challenges inherently lead to small sample sizes rarely exceeding N = 11 for wolves and N = 13 for dogs in contemporary studies where animals are hand-raised under identical conditions for species comparisons (Miklósi et al. 2003; Marshall-Pescini et al. 2017; Gácsi et al. 2005; Moretti et al. 2015; Range et al. 2015; Topál et al. 2005; Udell et al. 2012; Udell et al 2008). Hand-raising wolves and dogs under similar conditions is imperative, as behavioural development is highly influenced by environmental factors (Zimen 1987). Thus, because we heavily rely on these studies, with small sample sizes, to further increase our understanding of the domestication driven behavioural changes from wolf to dog, the importance of standardizing and reporting variation found across studies comparing wolves and dogs becomes critical. Furthermore, wolves used in studies to uncover the behavioural implications of domestication are predominantly North American wolves (Udell et al. 2008; Udell et al. 2012; Moretti et al. 2015; Range et al. 2015; Marshall-Pescini et al. 2017), and implementing standardized studies also on other sub-species of wolves might therefore add significant value to wolf-dog comparisons.

Here we examine the development of fear towards novelty in European grey wolves and dogs during the first six months of life, using standardized methods for both hand-raising, socializing (Klinghammer and Goodman 1987; Range and Virányi 2011; Udell et al. 2008) and testing (Marshall-Pescini et al. 2017; Moretti et al. 2015). We tested three litters of wolves (N = 13) and two litters of dogs (N = 12), hand-raised under identical conditions, at six, 10, 14, 18, 22 and 26 weeks of age, i.e. before sexual maturity, in repeated novel object tests. We used a new novel object in each of the six tests, choosing vastly different objects between tests to avoid the risk of habituation (van Oers et al. 2005; Noer et al. 2015). Novel objects were of different shape, size, colour and texture, and some objects included the element of sound and/or movement, similar to objects that have previously been used in novel object tests on dogs and wolves (Moretti et al. 2015; Marshall-Pescini et al. 2017). The novel object test is an established method to quantify fear and exploration of novelty and has been used on numerous species (Bremner-Harrison et al. 2004; Boogert et al. 2006; Mainwaring et al. 2011; Moretti et al. 2015; Marshall-Pescini et al. 2017). As is commonly applied in novel object tests, we used latency to approach and contact the novel object to quantify fear (Boissy 1995; Malmkvist and Hansen 2002; Meehan and Mench 2002; Ley et al. 2007; Moretti et al. 2015). Our longitudinal design allowed us to assess fear development and expression in juvenile wolves and dogs over an unprecedented period of time, and address our overall goal to test the hypothesis that domestication has altered fear responses in dogs compared to wolves. Based on studies reporting delayed onset of fear behaviour in domestic species (Martin and Fitzgerald 2005; Belyaev et al. 1985), including dogs and wolves (Coppinger and Coppinger 2001; Lord 2013), we expected wolves to express exaggerated fearfulness compared to dogs already at six to ten weeks of age by increased latency to approach the novel object and decreased exploratory behaviour. Furthermore, we predicted that domestication has lowered the interest of novelty in dogs (Brust and Guenther 2014; Moretti et al. 2015; Marshall-Pescini et al. 2017), and dogs therefore would express decreased interest in investigating and manipulating the novel object compared to wolves throughout the testing period.

## Materials and Methods

### Ethical statement

Daily care and all experiments were performed in accordance with relevant guidelines and regulations under national Swedish Law. The experimental protocols in this study were approved by the Ethical Committee in Uppsala, Sweden (approval number: C72/14). Facilities and daily care routines were approved by the Swedish National Board of Agriculture (approval number: 5.2.18-12309/13).

### Study animals

During 2014 – 2016 two litters of Alaskan huskies (N = 12) and three litters of European grey wolves (N = 13) were hand-raised and extensively socialized under similar conditions from the age of 10 days. This set-up was chosen to minimize environmental bias, including maternal effects, which is well-documented to affect the development of behavioural patterns (Wilsson and Sundgren 1998; Bray et al. 2017; Clark and Galef 1982). The Alaskan husky is a not a registered dog breed, but a type of dog specifically bred for dog sledding, consisting of a blend of registered dog breeds including Siberian Husky, Alaskan Malamute, Greenland Dog and various pointer breeds. Besides the issue of availability, Alaskan husky was our dog type of choice based on the morphological similarities with wolves (i.e. erect ears, similar size, long snouts etc.). This study was part of a bigger project to investigate domestication-driven changes in behavioural ontogeny in dogs, including social behaviour such as dominance. Thus, it was important to ensure that wolves and dogs had the same morphological basis providing them with equal opportunities to perform the same behavioural repertoires. Our choice of European grey wolves stands out as the majority of wolf-dog comparisons are based on North American wolves (Udell et al. 2008; Udell et al. 2012; Moretti et al. 2015; Range et al. 2015; Marshall-Pescini et al. 2017). The dog litter from 2014 consisted of five males and one female and the 2015 litter of three males and three females. The three wolf litters consisted of three females and two males in 2014, two males in 2015 and four males and two females in 2016.

Puppies were raised within litters and socialization involved 24-hour presence of human caregivers for the first two months. From two months of age, caregiver presence was decreased with a few hours a day until three months of age and then further decreased during every other night at four months of age. At six months of age, caregivers spent four to six hours with the puppies a day. All wolf and dog litters were kept separate, but reared under standardized conditions. From the age of 10 days to five weeks, puppies were reared in identical indoor rooms and here after given access to smaller roofed outdoor enclosures. After a week of habituation to the roofed outdoor enclosure, puppies were given access to a larger fenced grass enclosure at six weeks of age. Hereafter the puppies had free access to all three enclosures during the day and access to the indoor room and the roofed enclosure during the night. When the puppies where three months old they were moved to large outdoor enclosures (2,000 square meters), in which they remained for the rest of the study period. We started behavioural observations at 10 days of age and behavioural testing was initiated at six weeks of age. Testing procedures and exposure to the new environments were standardized over all three years. As required by national law, all hand-raisers were ethically certified and trained to handle animals. Furthermore, rules were implemented to assure that rearing was standardized across all caregivers. This included that puppies were never disciplined, trained or forced to have contact with their caregivers. From the age of eight weeks, puppies were gradually exposed to strangers through the fence with the support of one or more human caregivers.

### Experimental design

To investigate the ontogeny of fear expression in wolves and dogs, we designed a longitudinal experiment with novel object testing once a month starting at six weeks of age and ending at 26 weeks of age. The reason we chose to start testing at six weeks of age was to ensure that the puppies’ senses were fully developed (Lord 2013). Novel object tests were hereafter performed on a monthly basis at 10, 14, 18, 22 and 26 weeks of age using protocols similar to previous studies subjecting wolves and dogs to novel object tests (Marshall-Pescini et al. 2017; Moretti et al. 2015). To avoid environmental bias and disturbances by testing wolves and dogs in their out door home enclosures, we chose to conduct our tests in an indoor testing arena, which was familiar to both wolves and dogs. The equal familiarity among wolves and dogs with the test room also ensured that animals would focus on the novel object and not a novel environment (Moretti et al. 2015). In the test room (5×5 meters) a novel object was presented, placed opposite of where the puppy would enter the room, approximately four meters away from the door. This placement of the novel object ensured that puppies would actively have to approach the object to investigate and interact with it. Puppies were lead into the room by a caregiver, who quickly left the room and closed the door. The duration of a trial was 10 minutes and trials were always monitored. Some trials (N = 11, all wolves) were stopped prematurely to avoid destruction of the novel object. All test were filmed with two mounted GoPro cameras (model 3-4, GoPro Inc.) on opposite sides of the room.

### Novel objects

Due to the repeated exposure to novel objects in our experimental design, we chose vastly different objects between tests to avoid the risk of habituation (van Oers et al. 2005; Noer et al. 2015). We therefore chose novel objects of different shape, size, colour and texture, similar to objects that have previously been used for novel object tests on dogs and wolves (Moretti et al. 2015). Increasing the complexity of the novel object, such as adding sound or movement, has previously been used to avoid maturity and/or experience effects on habituation in novel object tests (Malmkvist et al. 2012). Thus, as a way of implementing complexity in later tests (week 22 and 26) we added movement and/or sound to the novel object, i.e. a mechanical dog and a moving bed sheet, respectively. Moving objects are well known to elicit fear responses (Boissy 1995) and mechanical toys have previously been used in novel object tests on wolves and/or dogs (Marshall-Pescini et al. 2017; Goddard and Beilharz 1984; Plutchik 1971; King et al. 2003). As we wished to test the response towards a fear eliciting stimuli in general, including social fear (Gray 1987), we opted to use a mirror as a novel object in week 14. While mirrors have previously been used in novel object tests to mimic a novel social context (Noer et al. 2015), we acknowledge that the use of a mirror to quantify fear responses might be considered controversial, and we therefore analysed our data both with and without the test at week 14 (see Statistical methods below).

According to procedures in previous novel object tests on wolves and dogs (Moretti et al. 2015), objects were handled as little as possible and always with freshly washed hands to avoid food smells transferring to the objects and possibly affecting the puppy’s behaviour towards the object. Novel objects chosen were at six weeks: a rolled up mattress, 10 weeks: a wheelbarrow (up-side down), 14 weeks: a mirror mounted to the wall, 18 weeks: a stuffed wolverine toy, 22 weeks: a moving mechanical dog and 24 weeks: a moving bed sheet (attached to a string).

### Behavioural scoring

As reported in other studies quantifying fear in dogs using novel objects (Stellato et al. 2017), the occurrence of subtle behaviours such as auto-grooming, vocalization, tail wagging and yawning was limited and we therefore chose to not include these behaviours in our analyses. The same was true for startle responses and piloerection, which were rarely expressed across tests. Differences in body posture are sometimes used as an indication of fear expression in dogs (Stellato et al. 2017; King et al. 2003). However, dogs can express altered body posture in neutral test conditions (i.e. no novel object present, (Stellato et al. 2017)). Thus, though dogs and wolves in our study were tested in a familiar room, we cannot rule out that confinement in an isolated room did not affect individuals differently. Therefore, to avoid potential bias by assessing body postures across individuals in two different species, we chose to use avoidance and approach (i.e. latency) behaviours related exclusively to the novel object to quantify fear. Avoidance behaviour and latency to approach a novel object is commonly applied to quantify fearfulness in various animal species (Boissy 1995; Malmkvist and Hansen 2002; Meehan and Mench 2002), including dogs and wolves (Ley et al. 2007; Moretti et al. 2015).

Behavioural scoring was carried out using the software BORIS v. 5.1.3. (Friard and Gamba 2016) based on an ethogram (Table 1). We chose our behavioural categories based on clear, non-overlapping segregation between behaviours directed at the novel object and behaviours not directed at the novel object, with prioritization of behaviours directed at the novel object. For instance, if the puppy was looking at the novel object while moving around the test room this was scored as *looking at novel object* and not *active behaviour*. We also attempted to graduate the behaviours directed at the novel object based on the puppies’ distance from the novel object. For example, we differentiated between the categories of *investigating novel object* and *looking at novel object*, based on how close the puppy was to the novel object (Table 1). Behaviours were logged in a non-overlapping way as durations, i.e. seconds (Table S1, S2 and S3). Similar to previous studies (Moretti et al. 2015), *latency to approach the novel object* was measured as the duration from test start to the time the puppy came within 1 meters distance of the novel object, and *latency to make contact with the novel object* was measured as the time lag to make physical contact with the novel object for the first time *after* the novel object had been approached within a distance of 1 meter. Based on cross coding, reliability of the behavioural scoring was calculated using Cohen’s kappa and was considered good with a value of 87.4%.

**Table 1.** Ethogram. Behaviours scored during novel object tests. Behaviours were scored in a non-overlapping way, with prioritization of behaviours related to the novel object. Latency times were measured regardless of the behaviour performed

| Behaviour | Description |
| --- | --- |
| Active behaviour | Moving around in, or interacting with, the test room with no attention to the novel object |
| Investigating novel object | Sniffing novel object or looking novel object form less than 1 meter |
| Latency to approach novel object | Time delay to approach the novel object with less than 1 meter |
| Latency make contact with novel object | Time delay to physically touch the novel object (sniffing) after having approached the object within a distance of less than 1 meter |
| Looking at novel object | Looking at novel object from a distance of more than 1 meter |
| Manipulating novel object | Pawing, nosing, scratching, biting, carrying, standing on novel object |
| Passive behaviour | Standing, sitting or lying passively with no attention to the novel object or the test room, including by the door |

### Statistical methods

We tested for the effect of species in each behaviour by fitting linear mixed models, with either latency or the time spent on a behaviour as the dependent variable. The fixed effects of interest were species, age, their interaction and sex. Additionally, for the models of time spent, we controlled for variation in the duration of each trial by including duration as a covariate. To account for the repeated measures of individuals and the non-independence of individuals from the same litter, we included random intercepts for both factors. The full model in lme4 syntax: y ~ species * age + sex + duration + (1|individual) + (1|litter). Models were then reduced by backwards model selection using AIC (cut-off ΔAIC > 2, Table S4), where the parameters for species, duration and the random effects were always maintained. Both latencies were log_10_ transformed, and the time spent looking, investigating and manipulating the novel object were log transformed after adding 1, in order to fulfil the assumption of normality in the model residuals. We centred the age variable to aid interpretation of the species effect in case of an interaction. When the interaction was retained in the model, we additionally fitted a model where age was a discrete variable, and used that to perform post-hoc tests for species differences at each age (Table S5 and S6). All p-values were obtained using Satterwaithe’s approximation of denominator degrees of freedom. Post-hoc p-values were corrected for multiple comparisons using the Holm method.

Because there were cases where the total duration of the test was less than 10 minutes, the total test duration was included as a covariate in our models. All but four puppies (dogs: N = 2, wolves: N = 2, Table S1) approached the novel object within a distance of 1 meter, and we assigned the total test time as latency to approach for the four puppies that did not approach the novel object. In eight cases (dogs: N = 6, wolves: N = 2) puppies did not make contact with the novel object (Table S1). Because it is inherently problematic for interpretation to assign a value to a non-occurring event, we took two different approaches to address this problem for latency to make contact to the novel object in our analyses. Following Marshall-Pescini et al. 2017, we performed the analysis for latency to make contact with the novel object using missing values for the eight cases were contact was not made (Table 2). We then repeated the analysis, using the lag time from the novel object was approached to the test ended as a measurement for non-occurring contact with the novel object. While the latter approach resulted in an overall species difference in latency to make contact with the novel object (Table S7 and S8), this difference was only significant at 22 weeks and this effect disappeared upon adjusting p-values for multiple comparisons (Holm method, Table S9). Therefore, the results from these two different approaches to analyse latency to make contact to the novel object were qualitatively the same, and we present results from the analyses using missing values for the eight cases were puppies did not make contact with the novel object below (Table 2).

**Table 2.** Model summary. Results for the best fitted model of repeated measures, with dogs as the reference, on 1) Latency to approach the novel object, 2) Latency to make contact with the novel object, 3) Looking at novel object (NO), 4) Investigating novel object, 5) Manipulating novel object, 6) Active behaviour and 7) Passive behaviour. Estimate, standard error, degrees of freedom, t-value and p-value are given. Significant p-values are marked in bold italic.

| *Behaviour* | *Term* | *Estimate* | *Std. error* | *df* | *t* | *p* |
| --- | --- | --- | --- | --- | --- | --- |
| **Latency, approach** | *(Intercept)* | 0.875 | 0.133 | 2.240 | 6.576 | ***0.017*** |
|  | *species* | 0.312 | 0.180 | 2.568 | 1.737 | 0.196 |
|  | *age* | -0.055 | 0.011 | 118.419 | -5.067 | ***<0.0001*** |
|  | *species:age* | 0.036 | 0.015 | 120.046 | 2.350 | ***0.020*** |
| **Latency, contact** | *(Intercept)* | 0.253 | 0.094 | 1.821 | 2.704 | 0.126 |
|  | *species* | 0.246 | 0.129 | 2.164 | 1.913 | 0.186 |
| **Looking at NO** | *(Intercept)* | 2.136 | 0.827 | 94.332 | 2.581 | ***0.011*** |
|  | *species* | 0.037 | 0.354 | 2.939 | 0.104 | 0.924 |
|  | *age* | 0.121 | 0.021 | 117.899 | 5.848 | ***<0.0001*** |
|  | *duration* | 0.001 | 0.001 | 124.417 | 0.426 | 0.671 |
|  | *species:age* | -0.064 | 0.031 | 120.667 | -2.058 | ***0.042*** |
| **Investigating NO** | *(Intercept)* | 0.416 | 0.717 | 126.889 | 0.580 | 0.563 |
|  | *species* | 0.141 | 0.234 | 3.493 | 0.601 | 0.585 |
|  | *age* | -0.119 | 0.019 | 138.727 | -6.384 | ***<0.0001*** |
|  | *duration* | 0.004 | 0.001 | 140.535 | 3.834 | ***0.0002*** |
|  | *species:age* | 0.056 | 0.028 | 139.315 | 1.994 | ***0.048*** |
| **Manipulating NO** | *(Intercept)* | 2.812 | 1.078 | 90.449 | 2.610 | ***0.011*** |
|  | *species* | 0.494 | 0.494 | 2.915 | 1.000 | 0.393 |
|  | *duration* | -0.001 | 0.002 | 141.262 | -0.786 | 0.433 |
| **Active behaviour** | *(Intercept)* | -145.265 | 57.693 | 97.323 | -2.518 | ***0.013*** |
|  | *species* | 102.697 | 24.105 | 2.977 | 4.260 | ***0.024*** |
|  | *age* | 2.587 | 1.176 | 122.299 | 2.200 | ***0.030*** |
|  | *duration* | 0.598 | 0.090 | 127.554 | 6.644 | ***<0.0001*** |
| **Passive behaviour** | *(Intercept)* | 152.645 | 54.263 | 25.688 | 2.813 | ***0.009*** |
|  | *species* | -91.478 | 39.977 | 3.003 | -2.288 | 0.106 |
|  | *age* | -4.106 | 0.962 | 121.459 | -4.268 | ***<0.0001*** |
|  | *duration* | 0.149 | 0.074 | 127.572 | 2.029 | ***0.045*** |

An alternative way of handling the latency measures is to conduct a survival analyses, since the trials with no observed latency can be handled as censored data. We therefore performed a survival analyses on latency to approach and latency to contact and found that the results were qualitatively the same as the main analyses (Table S10 and S11). We therefore present the main analyses with the transformed latency measures in the Results section.

To investigate whether the use of a mirror as a novel object in week 14 affected our conclusions, we chose to analyse our data with and without the test at 14 weeks. Upon excluding week 14 from our analyses we found, that while interaction terms for investigating and looking at the novel object disappeared, the results were overall similar to analyses including week 14 (Tables S12 and S13, Figures S1 and S2). Importantly, the exclusion of week 14 did not affect our main conclusion and we have therefore presented our results below based on our complete data set including week 14.

All statistical analyses were performed in R (v3.4.3, R Core Team 2016), with mixed effects models fitted using *lme4* v. 1.1-15 (Bates et al. 2015), survival analysis using *coxme* (Therneau 2018), Satterwaithe’s approximation from *lmerTest* v. 2.0-36 (Kuznetsova et al. 2017) and post-hoc testing using *emmeans* v. 1.1.2 (Lenth 2016).

## Results

### Latency measures

We found that wolves and dogs developed differently in latency to approach the novel object within 1 meter, where dogs expressed a larger reduction in latency with age compared to wolves (t = 2.35, df = 120.046, p = 0.02, Table 2, Figure 1 and 3). Dogs significantly decreased their latency with time, while wolves did not (see table S6 for slopes per species), resulting in dogs expressing significantly lowered latency to approach at 26 weeks compared to wolves (t = −3.131, df = 18.666, p = 0.006, p_adjusted_ = 0.034, Table S5). At younger ages we did not detect significant differences in latency to approach the novel object between dogs and wolves (Table S5).

**Figure 1.**
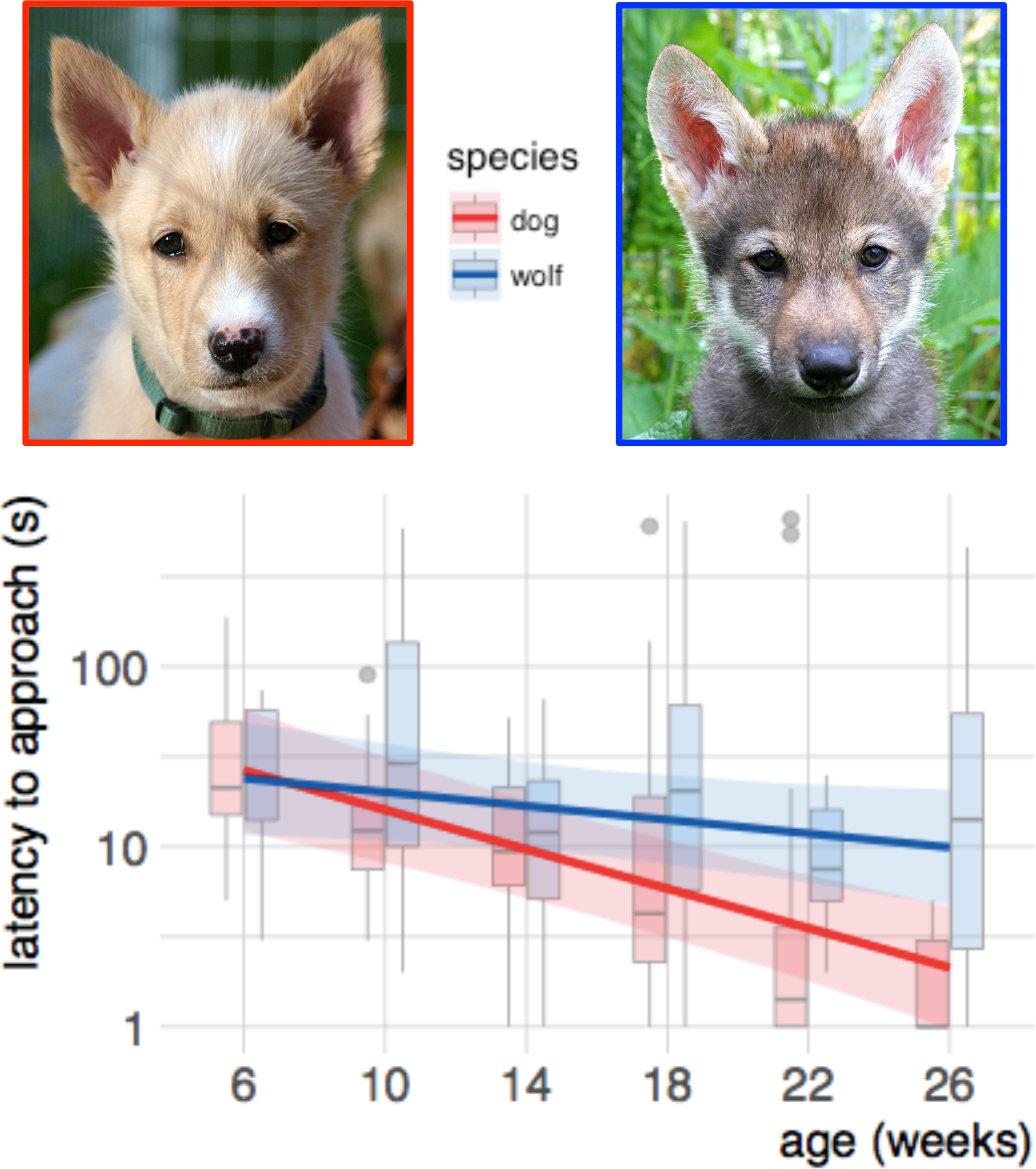
Dog – wolf comparisons, latency to approach. Boxplots shows behavioural scores during a novel object test, comparing dogs and wolves across age. Overlaid are the fits and confidence intervals from the best model, selected by AIC. Boxes indicate the quartiles, and the whiskers reach maximally 1.5 times the interquartile range. Values beyond that are shown as points. An a log(x) scale) was used. Photos: Christina Hansen Wheat

For the latency to make contact with the novel object, we found no differences in wolves and dogs (t = 1.931, df = 2.16, p = 0.186, Table 2, Figure 2a and 3), neither did we find evidence of sex differences in either species.

**Figure 2a-f.**
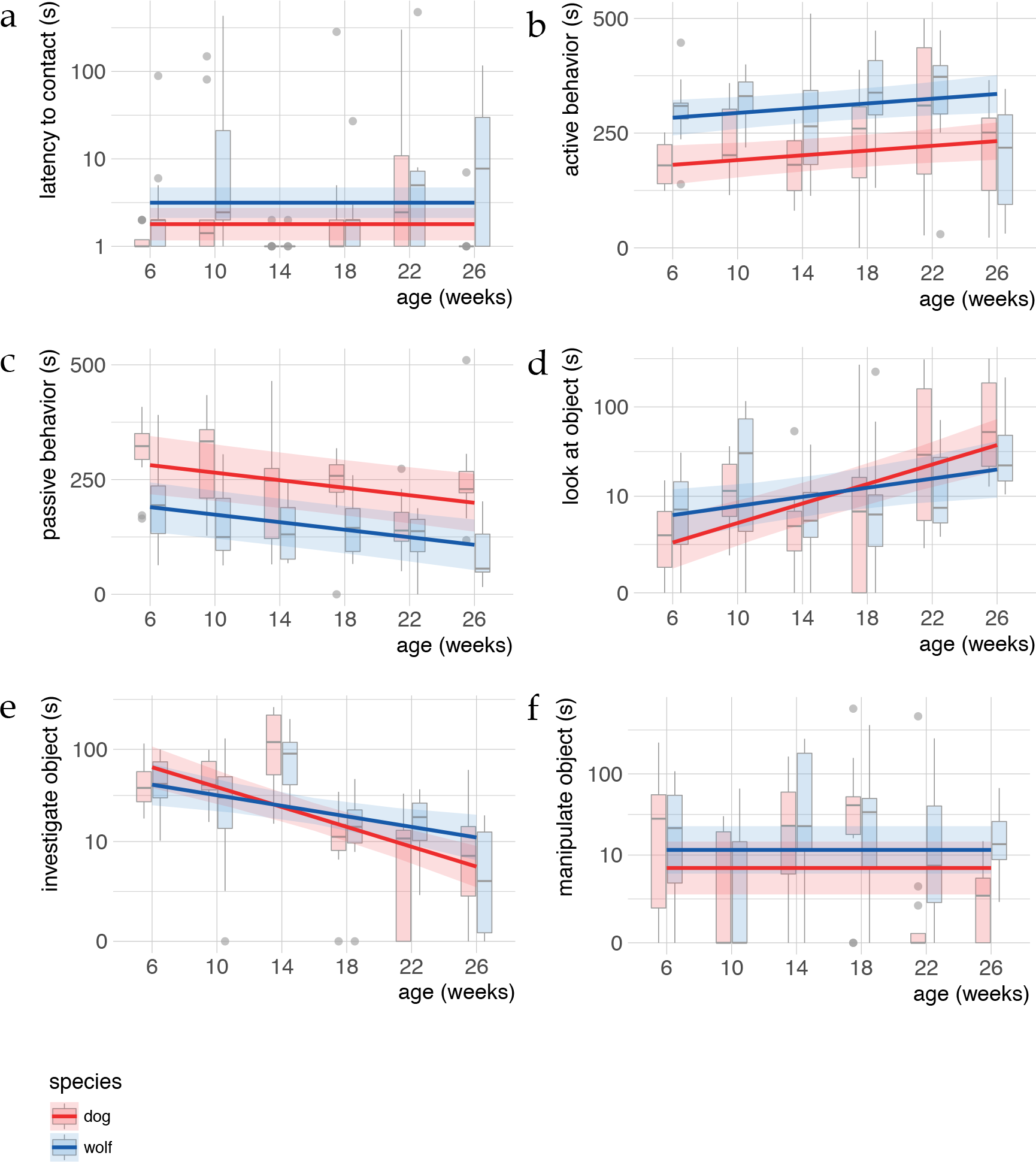
Dog – wolf comparisons. Boxplots shows behavioural scores during a novel object test, comparing dogs and wolves across age. Overlaid are the fits and confidence intervals from the best model, selected by AIC. Boxes indicate the quartiles, and the whiskers reach maximally 1.5 times the interquartile range. Values beyond that are shown as points. Note that b makes use of a log(x) scale, and panels d, e and f use log(x + 1).

**Figure 3.**
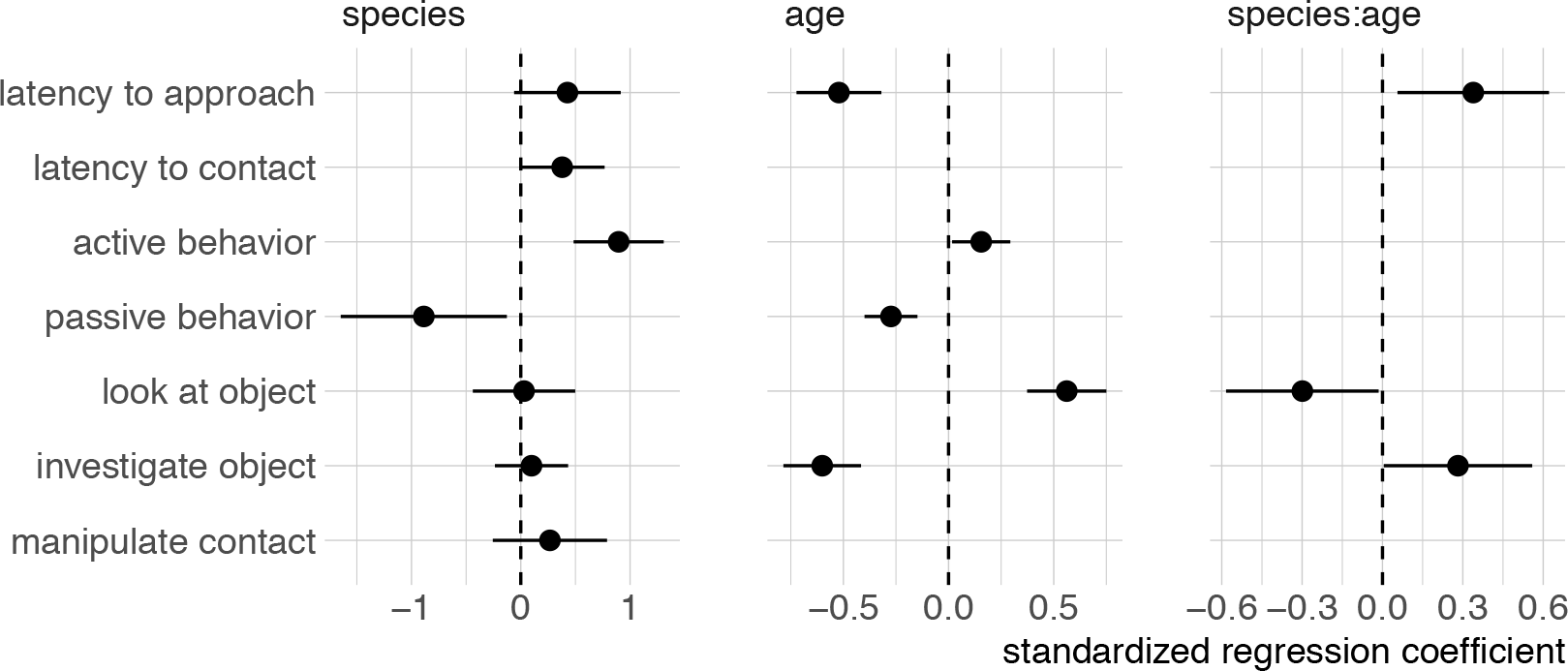
Standardized regression coefficients. Standardized regression coefficients for the best model for each behaviour, selected by AIC. Ranges indicate confidence intervals, computed using the likelihood profile. Missing estimates indicate that the term was not included in the best model.

### Behaviours related to the novel object

We found that wolves and dogs developed differently in looking at the novel object from a distance (t = −2.058, df = 120.667, p = 0.042, Table 1, Figure 2d and 3), but no such differences were detected in the post hoc tests (Table S5). While both wolves and dogs increased their time spent looking at the novel object from a distance with age (t = 5.848, df = 117.899, p <0.001, Table 2, Figure 2d), dogs expressed a stronger effect of age than wolves (Figure 2d, Table S6).

Wolves and dogs also showed different developmental trajectories for the time spent investigating the novel object (t = 1.994, df = 139.315, p = 0.048, Table 2, Figure 2e and 3). Post-hoc tests revealed that wolves investigated the novel object for longer at 22 weeks than dogs (t = −2.831, df = 28.029, p = 0.008, p_adjusted_ = 0.051, Figure 2e, Table S5). The significant interaction between species and age in investigating the novel object again consisted of stronger effect of age in dogs than in wolves (Figure 1f, Table S6), but with an overall decrease with age in both species (t = −6.384, df = 138.727, p <0.001, Table 2, Figure 2e). Wolves and dogs developed similarly in time spent manipulating the novel object (Table 2, Figure 2f and 3). There was no evidence of sex differences in either species.

### Behaviours not related to the novel object

We found that both species increased time spent on active behaviour with age (t = 2.2, df = 122.362, p = 0.03, Table 1, Figure 2b), with wolves expressing higher levels of activity than dogs (t = 4.26, df = 2.977, p = 0.024, Table 2, Figure 2b, Table S5). Passive behaviour decreased with age in both wolves and dogs (t = −4.268, df = 121.140, p <0.001, Table 2, Figure 2c), and while dogs appeared more passive than wolves the species differences was not significant. We found no evidence of sex differences.

## Discussion

Decreased expression of fear is considered a key behavioural alteration in domesticated animals, and it has further been suggested that domestication drives altered developmental rates delaying the initial onset of fear response (Belyaev et al. 1985). However, few studies have actually tested this experimentally and for wolves and dogs specifically, it remains unclear if and how a developmental shift during early ontogeny affects the continued development and expression of fear in either species. Here we present the first extended examination of the development of fear behaviour within the juvenile period in wolves and dogs. Contrary to expectations, we found no evidence in support of an increase in fearfulness in wolves with age or a delayed onset of fear response in dogs compared to wolves during early stages of development. Instead we found that dogs strongly reduced their fear response to a novel object in the period between six and 26 weeks of age. Critically, wolves did not differ in their fear response towards novelty over time, and the detected species difference was caused solely by a progressive reduced fear response in dogs. Furthermore, dogs and wolves did on average not differ in their interaction with the novel object. Together our results suggest that species differences in fear of novelty between wolves and dogs are not caused by a domestication driven shift in the first onset of fear response. Instead, we suggest that a loss of sensitivity towards novelty with increasing age in dogs causes the difference in fear expression towards novelty in wolves and dogs.

Fearfulness has previously been quantified by the latency to approach and explore novelty, and novel stimuli such as objects, arenas and people have been used to detect the timing of the initial onset of fear response in both wolves and dogs (Scott and Marston 1950; Freedman et al. 1961; Scott and Fuller 1965; Belyaev et al. 1985; Lord 2013; Morrow et al. 2015). However, while there is a general expectation that domestication has caused a delay in the sensitive period in dogs, resulting in later onset of fear behaviour compared to wolves (Scott and Fuller 1965; Fox 1970; Zimen 1987; Coppinger and Coppinger 2001; Lord 2013), we detected no species differences in fear expression during early development. This finding is in agreement with a recent study comparing exploration of novelty in six and eight weeks old wolves and dogs, which found no species differences in in fear behaviours or the latency to make contact with a novel object (Marshall-Pescini et al. 2017). Yet, it has been reported that adult wolves express increased latency to make contact to a novel object compared to dogs (Moretti et al. 2015), thereby suggesting that species differences in fear expression might arise later in development than previously thought. Thus, our finding that a species difference in latency to approach a novel object occurred at 26 weeks of age represents the first indication of when a quantifiable difference in fear towards novelty arises in wolves and dogs. We do, however, caution against an overly strong confidence in the exact timing of a species difference occurring at 26 weeks of age as it is possible that species differences emerge in the weeks prior, but that the current sample size is insufficient for earlier detection. Nonetheless, it is clear that a difference between species progressively develops towards the later end of the time period measured, and that we have captured the transition from equal expression of fear towards novelty in wolves and dogs to a clear species difference at 26 weeks of age. Importantly, the species difference in fear towards novelty did not occur because wolves became more fearful with age, as expected, but rather because dogs decreased their time to approach the novel object, which suggests that dogs, but not wolves lose their sensitivity towards novelty with age.

Upon subjecting individuals to repeated novel object tests, though objects differ between trials, there is a risk of habituation to novelty itself (Réale et al. 2007), and such a generalization of novelty *per se* can affect the potential to interpret fear responses from novel object tests. As such, one could speculate whether the decreased latency time to approach the novel object in dogs with age is a sign of habituation to the test situation itself. Furthermore, habituation to novelty itself can also be reflected in a decreased motivation to interact with a novel object (Zimmermann et al. 2001). Indeed, previous studies have demonstrated that while dogs and wolves show equal interest in approaching and exploring novel objects at 3-4 weeks of age (Gácsi et al. 2005), dogs seem to loose interest in investigating and interacting with novel objects with age (Moretti et al. 2015; Marshall-Pescini et al. 2017). While we do find an overall interaction between species and age in time spend investigating and looking at the novel object, wolves and dogs only differ in their investigation of the novel object at week 22 (the mechanical dog). Both wolves and dogs decrease the time they spend investigating the novel object with age, but increase the time they spend looking at the novel object with age, thereby suggesting that an interest in the novel object remains with age. Furthermore, the interaction effects seen in the investigation of and looking at the novel object does not extend to time spend on manipulating the novel object, in which neither wolves nor dogs differ with age. Thus, in our study wolves and dogs overall show equal interest in interacting with the novel object and we therefore find it unlikely that habituation to novelty in the dogs is driving our results.

The species difference we found in the latency to approach the novel object is not clearly reflected in differences in interaction with the same novel object. While fear of novelty was expressed immediately, through a delayed approach in wolves, once the novel object was approached this initial fearfulness appears to no longer affect behavioural responses in either species. This is reflected in wolves and dogs not differing in their latency to make contact with or interact with the novel object. While the latency to approach the novel object and the time spent being active and passive while in the test room showed consistent linear development over time in both wolves and dogs, the pattern in looking at, investigating and manipulating the novel object appeared variable across trials. This variability was most likely caused by the different novel objects that were used in the study, i.e. behaviours that are more closely related to the object itself show more variability across tests. It is possible that this increased variance may have prohibited detection of additional species differences in behavioural measures directly related to the novel object, such as increased exploration and manipulation of novel objects as reported in both juvenile and adult wolves compared to dogs (Marshall-Pescini et al. 2017; Moretti et al. 2015). Importantly, the linear development in latency to approach the novel object in both wolves dogs appeared to be less affected by the choice of novel object, indicating that latency to approach was more influenced by novelty itself.

Wolves develop physically faster than dogs (Frank and Frank 1982), and it has been suggested that wolves express increased activity at an earlier age than dogs due to this difference in developmental pace of motor patterns (Frank and Frank 1982; Marshall-Pescini et al. 2017). However, while we do find a species difference in how much time is spent on active behaviour during tests, this species difference is consistent across age and not restricted to early ontogeny alone. This indicates that wolves, on a general scale, are more active when in the test room than dogs. While it cannot be ruled out that active behaviour is affected by the presence of a novel object, it is a less likely explanation for our finding as we measured behaviours in a non-overlapping way with priority of behaviours related to the novel object. Thus, the measurement of activity does not include looking at, manipulating or approaching the novel object, but only time spent on active behaviour with no attention to the novel object. Instead the higher activity in wolves might reflect an increased reactivity of being separated from littermates and being confined in the test room compared to dogs.

Domestication has caused a general acceleration of sexual maturity in animals (Price 1999). Earlier sexual maturity in dogs (Morey 1994; Goodwin et al. 1997) could explain the steeper behavioural change observed in dogs compared to wolves across some of the behaviours related to the novel object in our study. However, while reproduction in wild living wolf packs is restricted to the breeding couple, it is currently unresolved if the lack of sexual activity in non-reproducing pack members is caused by delayed sexual maturity, behavioural suppression or restricted access to nutrition (Packard et al. 1985; Medjo and Mech 1976; Mech 1999). Nevertheless, it has been demonstrated that captive wolves removed from social constraints sexually mature as early as nine months of age (Medjo and Mech 1976). Thus, it is unclear if we should expect behavioural ontogeny to be affected by a shift in developmental pace caused by earlier sexual maturity when comparing wolves and dogs living in captive, non-reproductive groups. Our study was conducted before sexual maturity occurred in either wolves or dogs and as we found no effect of sex on the expression of behaviour, we suggest that the steeper development of some behaviours in dogs are instead related to the loss of sensitivity towards novelty.

Here we have compared behavioural development in wolves and dogs using standardized methods in both hand-raising, socialization (Klinghammer and Goodman 1987; Range and Virányi 2011; Udell et al. 2008) and testing (Marshall-Pescini et al. 2017; Moretti et al. 2015), thereby making our study comparable to some of the previous findings on fear development in the two species. Subsequently, our reporting of previously undetected variation in the development of fear expression is highly relevant for the on-going discussion of behavioural implications of domestication in dogs. In sum, our study shows that wolves and dogs do not differ in their fear towards novelty before late in the juvenile period. Importantly, the species difference does not occur because wolves become more fearful with age, but because dogs become less fearful with age. While it is possible that our novel results are reflected in our choice of European grey wolves, as the majority of studies on wolf dog comparison uses North American wolves, it emphasizes that different sub-species of wolves might reveal diverse behavioural variation. With more than 400 registered dog breeds in the world today (Lindblad-Toh et al. 2005), standardization of dog breeds used across studies comparing behaviours in wolves and dogs is less clear. Various dog breeds such as Poodle (Feddersen-Petersen 1991), Alaskan Malamute (Frank and Frank 1985) and German Shepherd, Siberian Husky, Alaskan Malamute, Czechoslovakian Wolfdog (Hansen Wheat et al. 2018) as well as mixed breeds (Range et al. 2015) have been used to uncover the behavioural implications of domestication from wolves, and with dogs being bred to fulfil highly specialized behavioural niches (Coppinger and Coppinger 2001; Mehrkam and Wynne 2014; Svartberg 2006), results will inevitably vary across studies (Morrow et al. 2015; Scott and Fuller 1965). However, detection of differences between wolves and dogs, no matter the breed of dog or subspecies of wolf, is of great importance to the continued discussion of the paradigm of domestication driven changes in behaviour. In conclusion, because of the small sample sizes inherently available in studies comparing behaviour in wolves and dogs, it is critical that continued, standardized studies on wolf dog comparisons are encouraged to further uncover the resolution in behavioural variation during domestication.

## Acknowledgements

We wish to thank the Department of Zoology, Stockholm University, for funding this study, our hand-raisers Patricia Berner, Anna Björk, Marjut Pokela, Linn Larsson, Charles Gent, Åsa Lycke, Erika Grasser, Joanna Schinner, Yrsa Andersson, Christoffer Sernert and Mija Jansson, and the staff at Tovetorp Zoological Research Station.

## Author contribution

CHW and HT designed the study, conducted the experiments and prepared data for analyses. WVDB and CHW planned how to analyse the data and WVDB analysed the data. CHW wrote the manuscript with input from HT and WVDB. All authors reviewed the manuscript prior to submission.

## References

Bates Douglas, Mächler M, Bolker B, Walker S. 2015. Fitting Linear Mixed-Effects Models Using Lme4. Journal of Statistical Software 67 (1). doi:10.18637/jss.v067.i01.

Belyaev, D K, Plyusnina I Z, Trut L N. 1985. Domestication in the Silver Fox (*Vulpes Fulvus Desm*): Changes in Physiological Boundaries of the Sensitive Period of Primary Socialization. Applied Animal Behaviour Science 13: 359–70. doi:10.1016/0168-1591(85)90015-2.

Bilkó Á, Altbäcker V. 2000. Regular Handling Early in the Nursing Period Eliminates Fear Responses Toward Human Beings in Wild and Domestic Rabbits. Developmental Psychobiology 36: 78–87. doi:10.1002/(SICI)1098-2302(200001)36:1

Boissy A. 1995. Fear and Fearfulness in Animals. The Quarterly Review of Biology 70: 165–91.

Boogert N J, Reader S M, Laland K N. 2006. The Relation Between Social Rank, Neophobia and Individual Learning in Starlings. Animal Behaviour 72: 1229–39. doi:10.1016/j.anbehav.2006.02.021.

Bray E E, Sammel M D, Cheney D L, Serpell J A, Seyfarth R M. 2017. Effects of Maternal Investment, Temperament, and Cognition on Guide Dog Success. PNAS 114 (34): 9128–33. doi:10.1073/pnas.1704303114.

Bremner-Harrison, S, Prodohl P A, Elwood R W. 2004. Behavioural Trait Assessment as a Release Criterion: Boldness Predicts Early Death in a Reintroduction Programme of Captive-Bred Swift Fox (*Vulpes Velox*). Animal Conservation 7 (3): 313–20. doi:10.1017/S1367943004001490.

Brust V, Guenther A. 2014. Domestication Effects on Behavioural Traits and Learning Performance: Comparing Wild Cavies to Guinea Pigs. Animal Cognition 18 (1): 99–109. doi:10.1016/0169-5347(94)90134-1.

Christensen J W, Zharkikh T, Ladewig J. 2008. Do Horses Generalise Between Objects During Habituation? Applied Animal Behaviour Science 114 (3-4): 509–20. doi:10.1016/j.applanim.2008.03.007.

Weidenmayer C P. 2009. Plasticity of Defensive Behavior and Fear in Early Development. Neuroscience & Biobehavioral Reviews 33 (3): 1447–57. doi:10.1016/j.neubiorev.2008.11.004.

Clark M M, Bennet G G Jr. 1982. Environmental Effects on the Ontogey of Exploratory and Escape Behaviors of Mongolian Gerbils. Developmental Psychobiology 15 (2): 121–29. doi:10.1002/dev.420150205.

Coppinger R, Coppinger L. 2001. Dogs: a Startling New Understanding of Canine Origin, Behavior & Evolution. The University of Chicago Press

Coppinger R, Glendinning J, Torop E, Matthay C, Sutherland M, Smith C. 1987. Degree of Behavioural Neoteny Differentiates Canid Polymorphs. Ethology 75: 89–108.

Crockford S J. 2002. Animal Domestication and Heterochronic Speciation: the Role of Thyroid Hormone. The Johns Hopkins University Press, 122–53.

Csatádi K, Kustos K, Eiben C, Bilkó Á, Altbäcker V. 2005. Even Minimal Human Contact Linked to Nursing Reduces Fear Responses Toward Humans in Rabbits. Applied Animal Behaviour Science 95 (1-2): 123–28. doi:10.1016/j.applanim.2005.05.002.

Dobney K, Larson G. 2006. Genetics and Animal Domestication: New Windows on an Elusive Process. Journal of Zoology 269 (0): 261–71. doi:10.1111/j.1469-7998.2006.00042.x.

Döring D, Roscher A, Scheipl F, Küchenhoff H, Erhard M H. 2009. Fear-Related Behaviour of Dogs in Veterinary Practice. The Veterinary Journal 182 (1). Elsevier Ltd: 38–43. doi:10.1016/j.tvjl.2008.05.006.

Driscoll C A, Macdonald D W, O’Brien S J. 2009. From Wild Animals to Domestic Pets, an Evolutionary View of Domestication. PNAS 106 (1): 9971–78.

Feddersen-Petersen D. 1991. The Ontogeny of Social Play and Agonistic Behaviour in Selected Canid Species. Bonn. Zool.Beitr. 2 (42): 97–114.

Fentress J C. 1967. Observations on the Behavioural Developement of a Hand-Reared Male Timber Wolf. American Zoologist 7 (2): 339–51.

Fox M W. 1970. A Comparative Study of the Development of Facial Expressions in Canids; Wolf, Coyote and Foxes. Behaviour 36 (1/2): 49–73.

Fox M W. 1972. Socio-Ecological Implications of Individual Differences in Wolf Litters : a Developmental and Evolutionary Perspective. Behaviour 41 (3): 298–313. doi:10.1163/156853972X00077.

Frank H, Frank M G. 1982. Comparison of Problem-Solving Performance in Six-Week-Old Wolves and Dogs. Animal Cognition 30: 95–98.

Frank H, Frank M G. 1985. Comparative Manipulation-Test Performance in Ten-Week-Old Wolves (Canis Lupus) and Alaskan Malamutes (Canis Familiaris): a Piagetian Interpretation. Journal of Comparative Psychology 99 (3): 266–74.

Freedman D G, King J A, Elliot O. 1961. Critical Period in the Social Development of Dogs. Science 31 (January): 1016–62.

Friard O, Gamba M. 2016. BORIS: a Free, Versatile Open-Source Event-Logging Software for Video/Audio Coding and Live Observations. Edited by Richard Fitzjohn. Methods in Ecology and Evolution 7 (11): 1325–30. doi:10.1006/anbe.1993.1127.

Gariépy J-L, Bauer D J, Cairns R B. 2001. Selective Breeding for Differential Aggression in Mice Provides Evidence for Heterochrony in Social Behaviours. Anima lBehaviour 61 (5): 933–47. doi:10.1006/anbe.2000.1700.

Gácsi M, Győri B, Miklósi Á, Virányi Z, Kubinyi E, Topál J, Csányi V. 2005. Species-Specific Differences and Similarities in the Behavior of Hand-Raised Dog and Wolf Pups in Social Situations with Humans. Developmental Psychobiology 47 (2): 111–22. doi:10.1002/dev.20082.

Goddard, M E, Beilharz R G. 1984. A Factor Analysis of Fearfulness in Potential Guide Dogs. Applied Animal Behaviour Science 12: 253–65.

Goodwin D, Bradshaw J W S, Wickens S M. 1997. Paedomorphosis Affects Agonistic Visual Signals of Domestic Dogs. Animal Behaviour 53: 297–304.

Gray J A. 1987. The Psychology of Fear and Stress. CUP Archive.

Griffin A S. 2004. Social Learning About Predators: a Review and Prospectus. Learning & Behavior 32 (1): 131–40. doi:10.3758/BF03196014.

Hansen Wheat C, Fitzpatrick J, Tapper I, Temrin H. 2018. Wolf (*Canis Lupus*) Hybrids Highlight the Importance of Human-Directed Play Behavior During Domestication of Dogs (*Canis Familiaris*). Journal of Comparative Psychology, 32 (4), 373–381. doi:10.1037/com0000119.

Hemsworth P H, Price E O, Borgwardt R. 1996. Behavioural Responses of Domestic Pigs and Cattle to Humans and Novel Stimuli. Applied Animal Behaviour Science 50: 43–56.

Jones R B, Waddington D. 1992. Modification of Fear in Domestic Chicks, *Gallus Gallus Domesticus*, via Regular Handling and Early Environmental Enrichment. Animal Behaviour, Short Communications 43: 1021–33.

King T, Hemsworth P H, Coleman G J. 2003. Fear of Novel and Startling Stimuli in Domestic Dogs. Applied Animal Behaviour Science 188 (1): 45–64. doi:10.1016/j.applanim.2016.12.007.

Klinghammer E, Goodman P A. 1987. Socialization and Management of Wolves in Captivity. In Man and Wolf, edited by Harry Frank. Dordrecht, Netherlands: Dr W Junk Publishers.

Kruska D. 1988. Mamalian Domestication and Its Effect on Brain Structure and Behaviour. Intelligence and Evolutionary Biology 17: 211–50.

Kuznetsova A, Brockhoff P B, Christensen R H B. 2017. lmerTestPackage: Tests in Linear Mixed Effects Models. Journal of Statistical Software 82 (13). doi:10.18637/jss.v082.i13.

Leiner L, Fendt M. 2011. Behavioural Fear and Heart Rate Responses of Horses After Exposure to Novel Objects: Effects of Habituation. Applied Animal Behaviour Science 131 (3-4). Elsevier B.V.: 104–9. doi:10.1016/j.applanim.2011.02.004.

Lenth R V. 2016. Least-Squares Means: the RPackage Lsmeans. Journal of Statistical Software 69 (1). doi:10.18637/jss.v069.i01.

Ley J, Coleman G J, Holmes R, Hemsworth P H. 2007. Assessing Fear of Novel and Startling Stimuli in Domestic Dogs. Applied Animal Behaviour Science 104 (1-2): 71–84. doi:10.1016/j.applanim.2006.03.021.

Lindblad-Toh K, Wade C M, Mikkelsen T S, Karlsson E K, Jaffe D B, Kamal M, Clamp M, et al. 2005. Genome Sequence, Comparative Analysis and Haplotype Structure of the Domestic Dog. Nature 438 (7069): 803–19. doi:10.1038/nature04338.

Lord K. 2013. A Comparison of the Sensory Development of Wolves (*Canis Lupus Lupus*) and Dogs (*Canis Lupus Familiaris*). Ethology 119 (2): 110–20. doi:10.1016/0304-3762(83)90109-8.

Mainwaring M C, Beal J L, Hartley I R. 2011. Zebra Finches Are Bolder in an Asocial, Rather Than Social, Context. Behavioural Processes 87 (2): 171–75. doi:10.1016/j.beproc.2011.03.005.

Malmkvist J, Hansen S W. 2002. Generalization of Fear in Farm Mink, Mustela Vison, Genetically Selected for Behaviour Towards Humans. Animal Behaviour 64 (3): 487–501. doi:10.1006/anbe.2002.3058.

Malmkvist J, Poulsen J M, Luthersson N, Palme R, Christensen J W, Søndergaard E. 2012. Behaviour and Stress Responses in Horses with Gastric Ulceration. Applied Animal Behaviour Science 142 (3-4): 160–67. doi:10.1016/j.applanim.2012.10.002.

Marshall-Pescini S, Virányi Z, Kubinyi E, Range F. 2017. Motivational Factors Underlying Problem Solving: Comparing Wolf and Dog Puppies’ Explorative and Neophobic Behaviors at 5, 6, and 8 Weeks of Age. Frontiers in Psychology 8: e20231. doi:10.1093/icb/7.2.357.

Martin J T. 1978. Embryonic Pituitary Adrenal Axis, Behavior Development and Domestication in Birds. American Zoologist 18: 489–99.

Martin L B II, Fitzgerald L. 2005. A Taste for Novelty in Invading House Sparrows, Passer Domesticus. Behavioral Ecology 16 (4): 702–7. doi:10.1006/anbe.2000.1725.

Mech D L. 1999. Alpha Status, Dominance, and Division of Labor in Wolf Packs. Can. J. Zool. 77: 1196–1203.

Medjo D C, Mech D L. 1976. Reproductive Activity in Nine-and Ten-Month_Old Wolves. Journal of Mammalogy 57 (May): 406–8. doi:10.2307/1379708.

Meehan C L, Mench J A. 2002. Environmental Enrichment Affects the Fear and Exploratory Responses to Novelty of Young Amazon Parrots. Applied Animal Behaviour Science 79: 75–88.

Mehrkam L R, Wynne C D L. 2014. Behavioral Differences Among Breeds of Domestic Dogs (*Canis Lupus Familiaris*): Current Status of the Science. Applied Animal Behaviour Science 155: 12–27. doi:10.1016/j.applanim.2014.03.005.

Miklósi Á, Kubinyi E, Topál J, Gácsi M, Virányi Z, Csányi V. 2003. A Simple Reason for a Big Difference. Current Biology 13 (9): 763–66. doi:10.1016/S0960-9822(03)00263-X.

Moretti L, Hentrup M, Kotrschal K, Range F. 2015. The Influence of Relationships on Neophobia and Exploration in Wolves and Dogs. Animal Behaviour 107: 159–73. doi:10.1016/j.anbehav.2015.06.008.

Morey D F. 1994. The Early Evolution of the Domestic Dog. American Scientist 82 (4): 336–47.

Morrow M, Ottobre J, Ottobre A, Neville P, St-Pierre N, Dreschel N, Pate J L. 2015. Breed-Dependent Differences in the Onset of Fear-Related Aviodance Behaviour in Puppies. Journal of Veterinary Behavior: Clinical Applications and Research 10: 286–94. doi:10.1016/j.jveb.2015.03.002.

Noer C L, Needham E K, Wiese A-S, Balsby T J S, Dabelsteen T. 2015. Context Matters: Multiple Novelty Tests Reveal Different Aspects of Shyness-Boldness in Farmed American Mink (*Neovison Vison*). PLoS ONE 10 (6): e0130474.doi:10.1371/journal.pone.0130474.s002.

Packard J M, Seal U S, Mech D L, Plotka E D. 1985. Causes of Reproductive Failure in Two Family Groups of Wolves (*Canis Lupus*). Zeitschrift Für Tierpsychologie 68 (1): 24–40. doi:10.1111/j.1439-0310.1985.tb00112.x.

Plutchik R. 1971. Individual and Breed Differences in Approach and Withdrawal in Dogs. Behaviour 40: 302–11.

Price E O. 2002. Animal Domestication and Behavior. CABI Publishing, CAB Int.

Price E O. 1999. Behavioral Development in Animals Undergoing Domestication. Applied Animal Behaviour Science 65: 245–71.

Range F, Ritter, Viranyi Z. 2015. Testing the Myth: Tolerant Dogs and Aggressive Wolves. Proceedings of the Royal Society of London B: Biological Sciences 282 (1807): 20150220–20. doi:10.1016/j.tics.2005.08.009.

Range F, Virányi Z. 2011. Development of Gaze Following Abilities in Wolves (Canis Lupus). PLoS ONE 6 (2): e16888. doi:10.1371/journal.pone.0016888.s002.

Réale D, Reader S M, Sol D, McDougall P T and Dingemanse N J. 2007. Integrating Animal Temperament Within Ecology and Evolution. Biological Reviews 82 (2): 291–318. doi:10.1016/S0091-6773(76)90901-9.

Scott J P. 1958. Critical Periods in the Development of Social Behavior in Puppies. Psychosomatic Medicine 20: 42–54. doi:10.1097/00006842-195801000-00005.

Scott J P. 1962. Critical Periods in Behavioural Development. Science 138: 949–58.

Scott J P, Fuller J L. 1965. Genetics and the Social Behavior of the Dog - the Classic Study. University of Chicago Press.

Scott J P, Marston M V. 1950. Critical Periods Affecting the Development of Normal and Mal-Adjustive Social Behavior of Puppies. The Pedagogical Seminary and Journal of Genetic Psychology 77 (1): 25–60. doi:10.1111/j.1749-6632.1950.tb27330.x.

Stellato A C, Flint H E, Widowski T M, Serpell J A, Niel L. 2017. Assessment of Fear-Related Behaviours Displayed by Companion Dogs (*Canis Familiaris*) in Response to Social and Non-Social Stimuli. Applied Animal Behaviour Science 188: 84–90. doi:10.1016/j.applanim.2016.12.007.

Svartberg K. 2006. Breed-Typical Behaviour in Dogs—Historical Remnants or Recent Constructs? Applied Animal Behaviour Science 96 (3-4): 293–313. doi:10.1016/j.applanim.2005.06.014.

Therneau T M. 2018. Mixed Effects Cox Models. Cran, May, 1–21. https://CRAN.R-project.org/package=coxme.

Topál J, Gácsi M, Miklósi Á, Virányi Z, Kubinyi E, Csányi V. 2005. Attachment to Humans: a Comparative Study on Hand-Reared Wolves and Differently Socialized Dog Puppies. Animal Behaviour 70 (6): 1367–75. doi:10.1016/j.anbehav.2005.03.025.

Trut L N, Plyusnina I Z, Oskina I N. 2004. An Experiment on Fox Domestication and Debatable Issues of Evolution of the Dog. Russian Journal of Genetics 40 (6): 644–55. doi:10.1023/B:RUGE.0000033312.92773.c1.

Trut L N. 1999. Early Canid Domestication: the Farm-Fox Experiment: Foxes Bred for Tamability in a 40-Year Experiment Exhibit Remarkable Transformations That Suggest an Interplay Between Behavioral Genetics and Development. American Scientist 87: 160–69.

Trut L N, Oskina I N, Kharlamova A. 2009. Animal Evolution During Domestication: the Domesticated Fox as a Model. Bioessays 31 (3): 349–60. doi:10.1002/bies.200800070.

Udell M A R, Spencer J M, Dorey N R, Wynne C D L. 2012. Human-Socialized Wolves Follow Diverse Human Gestures…and They May Not Be Alone. International Journal of Comparative Psychology 25: 97–177.

Udell M A R, Dorey N R, Wynne C D L. 2008 Wolves Outperform Dogs in Following Human Social Cues. Animal Behaviour 76 (6): 1767–73. doi:10.1016/j.anbehav.2008.07.028.

van Oers K, Klunder M, Drent P J. 2005. Context Dependence of Personalities: Risk-Taking Behavior in a Social and a Nonsocial Situation. Behavioral Ecology 16 (4): 716–23. doi:10.1016/0168-1591(95)00557-9.

Wilkins A S, Wrangham R W, Fitch W T. 2014. The ‘Domestication Syndrome’ in Mammals: a Unified Explanation Based on Neural Crest Cell Behavior and Genetics. Genetics 197 (3): 795–808. doi:10.1534/genetics.114.165423.

Wilsson E, Sundgren P-E. 1998. Behaviour Test for Eight-Week Old Puppies— Heritabilities of Tested Behaviour Traits and Its Correspondence to Later Behaviour. Applied Animal Behaviour Science 58: 151–62.

Wooply J H, Ginsburg B E. 1967. Wolf Socialization: a Study of Temperament in a Wild Social Species. American Zoologist 7 (2): 357–63.

Zeder M A. 2012. The Domestication of Animals. Journal of Anthropological Research 68: 161–90.

Zimen E. 1987. Ontogeny of Approach and Flight Behavior Towards Humans in Wolves, Poodles and Wolf-Poodle Hybrids. In Man and Wolf, edited by H Frank. Dr. W. Junk Publishers, Dordrecht.

Zimmerman A, Stauffacher M, Langhans W, Würbel H. 2001. Enrichment-Dependent Differences in Novelty Exploration in Rats Can Be Explained by Habituation. Behavioural Brain Research 121 (1-2): 11–20.

