## Supplementary materials for "Dogs, but not wolves, lose their sensitivity towards novelty with age"

Table S1. Behavioural scores, latencies……………………………………… 2

Table S2. Behavioural scores, novel object related………………………… 6

Table S3. Behavioural scores, non-novel object related…….……………….. 10

Table S4. Model selection....…………...……………………………………... 14

Table S5. Post hoc testing, contrasts ……………….……..………………… 15

Table S6. Post hoc testing, slopes……………….……….…………………... 16

Table S12. Model selection, excluding data for week 14……………….… 22

Table S13. . Model summary, excluding data for week 14...……………..... 23

Figure S1. Dog – wolf comparisons, excluding data for week 14……….. 24

Figure S2. Standardized regression coefficients, excl. week 14………… 25

**Table S1. Behavioural scores, latencies.** Latencies for approaching and making contact with the novel object and total test time for each of the six novel object tests. Latency to approach (Lat_app) is measured as the time in seconds from test start to the puppy comes within a 1 meter radius of the novel object. Latency to make contact (Lat_con) is measured as the time lag in seconds to make physical contact with the novel object after it has been approached. If a puppy never approached the novel object, the full test time was used. If a puppy never made contact to the novel object "NA" (Table 2) or lag time (parenthesis) from approach to total test time was used (Table S7).

| Individual | Sex | Species | Litter | Novel object | Lat_app | Lat_con | Total time |
|---|---|---|---|---|---|---|---|
| Bowie_6w | M | dog | 2014 | Rolled up mattress | 5 | 1 | 600.076 |
| Bowie_10w | M | dog | 2014 | Wheelbarrow | 90 | 1 | 587.139 |
| Bowie_14w | M | dog | 2014 | Mirror | 35 | 1 | 612.262 |
| Bowie_18w | M | dog | 2014 | Stuffed wolverine | 26 | 1 | 609.377 |
| Bowie_22w | M | dog | 2014 | Mechanical dog | 1 | 1 | 603.233 |
| Bowie_26w | M | dog | 2014 | Moving sheet | 1 | NA (575.78) | 576.78 |
| Cash_6w | M | dog | 2014 | Rolled up mattress | 46 | 2 | 601.049 |
| Cash_10w | M | dog | 2014 | Wheelbarrow | 54 | 1 | 627.571 |
| Cash_14w | M | dog | 2014 | Mirror | 12 | 1 | 574.068 |
| Cash_18w | M | dog | 2014 | Stuffed wolverine | 1 | 1 | 614.802 |
| Cash_22w | M | dog | 2014 | Mechanical dog | 1 | 301 | 607.46 |
| Cash_26w | M | dog | 2014 | Moving sheet | 1 | 1 | 609.681 |
| Jagger_6w | M | dog | 2014 | Rolled up mattress | 188 | 1 | 603.606 |
| Jagger_10w | M | dog | 2014 | Wheelbarrow | 32 | 1 | 604.013 |
| Jagger_14w | M | dog | 2014 | Mirror | 52 | 1 | 609.974 |
| Jagger_18w | M | dog | 2014 | Stuffed wolverine | 1 | 1 | 605.033 |
| Jagger_22w | M | dog | 2014 | Mechanical dog | 1 | NA (616.042) | 617.042 |
| Jagger_26w | M | dog | 2014 | Moving sheet | 1 | 1 | 605.032 |
| Janis_6w | F | dog | 2014 | Rolled up mattress | 60 | 1 | 604.807 |
| Janis_10w | F | dog | 2014 | Wheelbarrow | 10 | 1 | 608.57 |
| Janis_14w | F | dog | 2014 | Mirror | 4 | 1 | 622.112 |
| Janis_18w | F | dog | 2014 | Stuffed wolverine | 138 | 2 | 606.243 |
| Janis_22w | F | dog | 2014 | Mechanical dog | 658.976 | NA (0) | 658.976 |
| Janis_26w | F | dog | 2014 | Moving sheet | 1 | 1 | 594.822 |
| Lennon_6w | M | dog | 2014 | Rolled up mattress | 82 | 2 | 606.69 |
| Lennon_10w | M | dog | 2014 | Wheelbarrow | 15 | 149 | 610.654 |
| Lennon_14w | M | dog | 2014 | Mirror | 22 | 1 | 601.694 |
| Lennon_18w | M | dog | 2014 | Stuffed wolverine | 1 | 1 | 616.13 |
| Lennon_22w | M | dog | 2014 | Mechanical dog | 1 | 9 | 606.609 |
| Lennon_26w | M | dog | 2014 | Moving sheet | 1 | 1 | 594.675 |
| Marley_6w | M | dog | 2014 | Rolled up mattress | 30 | 1 | 604.525 |
| Marley_10w | M | dog | 2014 | Wheelbarrow | 3 | 1 | 623.797 |
| Marley_14w | M | dog | 2014 | Mirror | 3 | 1 | 613.112 |
| Marley_18w | M | dog | 2014 | Stuffed wolverine | 601.682 | NA (0) | 601.682 |
| Marley_22w | M | dog | 2014 | Mechanical dog | 1 | NA (605.481) | 606.481 |
| Marley_26w | M | dog | 2014 | Moving sheet | 1 | 1 | 603.068 |

*Table S1, continued*

| Individual | Sex | Species | Litter | Novel object | Lat_app | Lat_con | Total time |
|---|---|---|---|---|---|---|---|
| Björk_6w | F | wolf | 2014 | Rolled up mattress | 36 | 1 | 579.774 |
| Björk_10w | F | wolf | 2014 | Wheelbarrow | 3 | 1 | 595.852 |
| Björk_14w | F | wolf | 2014 | Mirror | 1 | 1 | 631.068 |
| Björk_18w | F | wolf | 2014 | Stuffed wolverine | 5 | 1 | 616.321 |
| Björk_22w | F | wolf | 2014 | Mechanical dog | 2 | 2 | 598.221 |
| Björk_26w | F | wolf | 2014 | Moving sheet | 60 | 1 | 609.076 |
| Iggy_6w | M | wolf | 2014 | Rolled up mattress | 6 | 2 | 600.857 |
| Iggy_10w | M | wolf | 2014 | Wheelbarrow | 28 | 1 | 610.047 |
| Iggy_14w | M | wolf | 2014 | Mirror | 13 | 1 | 615.548 |
| Iggy_18w | M | wolf | 2014 | Stuffed wolverine | 5 | 2 | 623.099 |
| Iggy_22w | M | wolf | 2014 | Mechanical dog | 16 | 477 | 638.666 |
| Iggy_26w | M | wolf | 2014 | Moving sheet | 1 | 28 | 414.633 |
| Joni_6w | F | wolf | 2014 | Rolled up mattress | 6 | 2 | 573.469 |
| Joni_10w | F | wolf | 2014 | Wheelbarrow | 236 | 3 | 632.978 |
| Joni_14w | F | wolf | 2014 | Mirror | 11 | 1 | 649.435 |
| Joni_18w | F | wolf | 2014 | Stuffed wolverine | 70 | 2 | 587.12 |
| Joni_22w | F | wolf | 2014 | Mechanical dog | 5 | 5 | 609.673 |
| Joni_26w | F | wolf | 2014 | Moving sheet | 460 | 68 | 612.939 |
| Lita_6w | F | wolf | 2014 | Rolled up mattress | 14 | 1 | 616.709 |
| Lita_10w | F | wolf | 2014 | Wheelbarrow | 5 | 2 | 623.776 |
| Lita_14w | F | wolf | 2014 | Mirror | 66 | 1 | 617.91 |
| Lita_18w | F | wolf | 2014 | Stuffed wolverine | 16 | 2 | 624.842 |
| Lita_22w | F | wolf | 2014 | Mechanical dog | 7 | 5 | 593.112 |
| Lita_26w | F | wolf | 2014 | Moving sheet | 2 | 1 | 238.123 |
| Ozzy_6w | M | wolf | 2014 | Rolled up mattress | 24 | 1 | 589.697 |
| Ozzy_10w | M | wolf | 2014 | Wheelbarrow | 10 | 2 | 619.074 |
| Ozzy_14w | M | wolf | 2014 | Mirror | 11 | 1 | 626.846 |
| Ozzy_18w | M | wolf | 2014 | Stuffed wolverine | 8 | 1 | 596.166 |
| Ozzy_22w | M | wolf | 2014 | Mechanical dog | 25 | 8 | 256.226 |
| Ozzy_26w | M | wolf | 2014 | Moving sheet | 4 | 36 | 630.475 |
| Billie_6w | F | dog | 2015 | Rolled up mattress | 16 | 1 | 618.307 |
| Billie_10w | F | dog | 2015 | Wheelbarrow | 17 | 2 | 606.824 |
| Billie_14w | F | dog | 2015 | Mirror | 8 | 2 | 622.631 |
| Billie_18w | F | dog | 2015 | Stuffed wolverine | 13 | 5 | 617.803 |
| Billie_22w | F | dog | 2015 | Mechanical dog | 542 | NA (77.881) | 619.881 |
| Billie_26w | F | dog | 2015 | Moving sheet | 1 | 1 | 624.415 |
| Ella_6w | F | dog | 2015 | Rolled up mattress | 15 | 1 | 617.097 |
| Ella_10w | F | dog | 2015 | Wheelbarrow | 15 | 2 | 608.453 |
| Ella_14w | F | dog | 2015 | Mirror | 1 | 1 | 624.353 |
| Ella_18w | F | dog | 2015 | Stuffed wolverine | 17 | 284 | 617.032 |
| Ella_22w | F | dog | 2015 | Mechanical dog | 21 | 19 | 608.397 |
| Ella_26w | F | dog | 2015 | Moving sheet | 3 | 1 | 605.549 |

*Table S1, continued*

| Individual | Sex | Species | Litter | Novel object | Lat_app | Lat_con | Total time |
|---|---|---|---|---|---|---|---|
| Muddy_6w | M | dog | 2015 | Rolled up mattress | 15 | 2 | 614.377 |
| Muddy_10w | M | dog | 2015 | Wheelbarrow | 9 | 1 | 625.693 |
| Muddy_14w | M | dog | 2015 | Mirror | 8 | 1 | 619.128 |
| Muddy_18w | M | dog | 2015 | Stuffed wolverine | 3 | 2 | 614.359 |
| Muddy_22w | M | dog | 2015 | Mechanical dog | 2 | 1 | 622.874 |
| Muddy_26w | M | dog | 2015 | Moving sheet | 3 | 1 | 600.184 |
| Red_6w | M | dog | 2015 | Rolled up mattress | 18 | 1 | 612.618 |
| Red_10w | M | dog | 2015 | Wheelbarrow | 8 | 2 | 631.432 |
| Red_14w | M | dog | 2015 | Mirror | 7 | 1 | 618.505 |
| Red_18w | M | dog | 2015 | Stuffed wolverine | 3 | 1 | 610.963 |
| Red_22w | M | dog | 2015 | Mechanical dog | 2 | 3 | 626.564 |
| Red_26w | M | dog | 2015 | Moving sheet | 1 | 1 | 605.679 |
| Simone_6w | F | dog | 2015 | Rolled up mattress | 25 | 1 | 621.930 |
| Simone_10w | F | dog | 2015 | Wheelbarrow | 5 | 2 | 605.543 |
| Simone_14w | F | dog | 2015 | Mirror | 21 | 1 | 609.914 |
| Simone_18w | F | dog | 2015 | Stuffed wolverine | 3 | 1 | 624.499 |
| Simone_22w | F | dog | 2015 | Mechanical dog | 1 | 1 | 615.770 |
| Simone_26w | F | dog | 2015 | Moving sheet | 5 | 1 | 610.149 |
| Skip_6w | M | dog | 2015 | Rolled up mattress | 7 | 1 | 610.773 |
| Skip_10w | M | dog | 2015 | Wheelbarrow | 6 | 81 | 631.961 |
| Skip_14w | M | dog | 2015 | Mirror | 11 | 1 | 621.581 |
| Skip_18w | M | dog | 2015 | Stuffed wolverine | 6 | 2 | 614.454 |
| Skip_22w | M | dog | 2015 | Mechanical dog | 2 | 2 | 617.542 |
| Skip_26w | M | dog | 2015 | Moving sheet | 3 | 7 | 610.066 |
| Flea_6w | M | wolf | 2015 | Rolled up mattress | 3 | 1 | 587.411 |
| Flea_10w | M | wolf | 2015 | Wheelbarrow | 172 | 40 | 598.813 |
| Flea_14w | M | wolf | 2015 | Mirror | 22 | 1 | 609.896 |
| Flea_18w | M | wolf | 2015 | Stuffed wolverine | 108 | 2 | 609.496 |
| Flea_22w | M | wolf | 2015 | Mechanical dog | 16 | 8 | 602.450 |
| Flea_26w | M | wolf | 2015 | Moving sheet | 25 | 2 | 95.161 |
| Hendrix_6w | M | wolf | 2015 | Rolled up mattress | 14 | 1 | 593.077 |
| Hendrix_10w | M | wolf | 2015 | Wheelbarrow | 2 | 2 | 587.873 |
| Hendrix_14w | M | wolf | 2015 | Mirror | 2 | 1 | 612.873 |
| Hendrix_18w | M | wolf | 2015 | Stuffed wolverine | 1 | 1 | 623.254 |
| Hendrix_22w | M | wolf | 2015 | Mechanical dog | 8 | 1 | 608.376 |
| Hendrix_26w | M | wolf | 2015 | Moving sheet | 2 | 1 | 488.468 |
| Elvis_6w | M | wolf | 2016 | Rolled up mattress | 67 | 2 | 612.777 |
| Elvis_10w | M | wolf | 2016 | Wheelbarrow | 29 | 433 | 626.783 |
| Elvis_14w | M | wolf | 2016 | Mirror | 27 | 2 | 637.202 |
| Elvis_18w | M | wolf | 2016 | Stuffed wolverine | 26 | 1 | 630.121 |
| Elvis_22w | M | wolf | 2016 | Mechanical dog | 2 | 1 | 604.293 |
| Elvis_26w | M | wolf | 2016 | Moving sheet | 54 | 20 | 128.490 |

*Table S1, continued*

| Individual | Sex | Species | Litter | Novel object | Lat_app | Lat_con | Total time |
|---|---|---|---|---|---|---|---|
| KD_6w | F | wolf | 2016 | Rolled up mattress | 74 | 5 | 609.205 |
| KD_10w | F | wolf | 2016 | Wheelbarrow | 136 | 5 | 637.961 |
| KD_14w | F | wolf | 2016 | Mirror | 48 | 1 | 652.284 |
| KD_18w | F | wolf | 2016 | Stuffed wolverine | 58 | 27 | 620.433 |
| KD_22w | F | wolf | 2016 | Mechanical dog | 7 | 7 | 613.584 |
| KD_26w | F | wolf | 2016 | Moving sheet | 3 | 117 | 481.627 |
| Lemmy_6w | M | wolf | 2016 | Rolled up mattress | 46 | 2 | 606.694 |
| Lemmy_10w | M | wolf | 2016 | Wheel barrel | 586.976 | NA (0) | 586.976 |
| Moby_6w | M | wolf | 2016 | Rolled up mattress | 64 | 1 | 617.363 |
| Moby_10w | M | wolf | 2016 | Wheelbarrow | 130 | 192 | 627.734 |
| Moby_14w | M | wolf | 2016 | Mirror | 18 | 1 | 630.257 |
| Moby_18w | M | wolf | 2016 | Stuffed wolverine | 641.919 | NA (0) | 641.919 |
| Moby_22w | M | wolf | 2016 | Mechanical dog | 15 | 5 | 602.346 |
| Moby_26w | M | wolf | 2016 | Moving sheet | 57 | 3 | 106.774 |
| PJ_6w | F | wolf | 2016 | Rolled up mattress | 57 | 89 | 580.513 |
| PJ_10w | F | wolf | 2016 | Wheelbarrow | 87 | 17 | 615.866 |
| PJ_14w | F | wolf | 2016 | Mirror | 7 | 1 | 616.296 |
| PJ_18w | F | wolf | 2016 | Stuffed wolverine | 6 | 1 | 603.243 |
| PJ_22w | F | wolf | 2016 | Mechanical dog | 5 | 1 | 430.614 |
| PJ_26w | F | wolf | 2016 | Moving sheet | 8 | 27 | 468.549 |
| Sting_6w | M | wolf | 2016 | Rolled up mattress | 15 | 6 | 608.304 |
| Sting_10w | M | wolf | 2016 | Wheelbarrow | 25 | 2 | 616.169 |
| Sting_14w | M | wolf | 2016 | Mirror | 1 | 1 | 629.972 |
| Sting_18w | M | wolf | 2016 | Stuffed wolverine | 31 | 3 | 616.396 |
| Sting_22w | M | wolf | 2016 | Mechanical dog | 20 | 1 | 600.794 |
| Sting_26w | M | wolf | 2016 | Moving sheet | 28 | 1 | 194.205 |

**Table S2. Behavioural scores, novel object related.** Scores (seconds) for behaviours related to the novel object for each of the six novel object tests.

| Individual | Sex | Species | Litter | Novel object | Invest_NO | Look_NO | Manip_NO |
|---|---|---|---|---|---|---|---|
| Bowie_6w | M | dog | 2014 | Rolled up mattress | 23.011 | | 41.449 |
| Bowie_10w | M | dog | 2014 | Wheelbarrow | 39.913 | 2.298 | 30.809 |
| Bowie_14w | M | dog | 2014 | Mirror | 55.28 | | |
| Bowie_18w | M | dog | 2014 | Stuffed wolverine | 9.95 | 17.013 | 45.067 |
| Bowie_22w | M | dog | 2014 | Mechanical dog | 33.833 | 26.865 | 3.672 |
| Bowie_26w | M | dog | 2014 | Moving sheet | | 49.256 | |
| Cash_6w | M | dog | 2014 | Rolled up mattress | 43.257 | 3.545 | |
| Cash_10w | M | dog | 2014 | Wheelbarrow | 29.264 | 21.581 | 19.779 |
| Cash_14w | M | dog | 2014 | Mirror | 49.439 | 1.296 | 35.033 |
| Cash_18w | M | dog | 2014 | Stuffed wolverine | 7.741 | | 44.245 |
| Cash_22w | M | dog | 2014 | Mechanical dog | 25.863 | 3.595 | |
| Cash_26w | M | dog | 2014 | Moving sheet | 30.219 | 25.534 | |
| Jagger_6w | M | dog | 2014 | Rolled up mattress | 18.154 | 2.828 | 46.103 |
| Jagger_10w | M | dog | 2014 | Wheelbarrow | 33.536 | 17.959 | |
| Jagger_14w | M | dog | 2014 | Mirror | 232.6 | 5.374 | 15.608 |
| Jagger_18w | M | dog | 2014 | Stuffed wolverine | 7.937 | | 16.449 |
| Jagger_22w | M | dog | 2014 | Mechanical dog | | 90.557 | |
| Jagger_26w | M | dog | 2014 | Moving sheet | 5.998 | 59.871 | |
| Janis_6w | F | dog | 2014 | Rolled up mattress | 28.373 | 6.932 | 2.557 |
| Janis_10w | F | dog | 2014 | Wheelbarrow | 38.738 | 1.538 | |
| Janis_14w | F | dog | 2014 | Mirror | 72.683 | 3.309 | 8.708 |
| Janis_18w | F | dog | 2014 | Stuffed wolverine | 16.426 | 36.963 | 41.729 |
| Janis_22w | F | dog | 2014 | Mechanical dog | | 254.17 | |
| Janis_26w | F | dog | 2014 | Moving sheet | 60.585 | 156.475 | |
| Lennon_6w | M | dog | 2014 | Rolled up mattress | 25.589 | 3.598 | 19.737 |
| Lennon_10w | M | dog | 2014 | Wheelbarrow | 16.628 | 29.413 | 19.729 |
| Lennon_14w | M | dog | 2014 | Mirror | 15.913 | 54.314 | |
| Lennon_18w | M | dog | 2014 | Stuffed wolverine | 21.243 | 4.843 | 42.025 |
| Lennon_22w | M | dog | 2014 | Mechanical dog | 11.504 | 159.205 | |
| Lennon_26w | M | dog | 2014 | Moving sheet | 11.572 | 22.247 | 3.574 |
| Marley_6w | M | dog | 2014 | Rolled up mattress | 28.422 | 15.309 | 7.957 |
| Marley_10w | M | dog | 2014 | Wheelbarrow | 44.044 | 7.647 | |
| Marley_14w | M | dog | 2014 | Mirror | 39.131 | 3.065 | 1.011 |
| Marley_18w | M | dog | 2014 | Stuffed wolverine | | 287.018 | |
| Marley_22w | M | dog | 2014 | Mechanical dog | | 327.468 | |
| Marley_26w | M | dog | 2014 | Moving sheet | 7.697 | 333.993 | 1.54 |

*Table S2, continued*

| Individual | Sex | Species | Litter | Novel object | Invest_NO | Look_NO | Manip_NO |
|---|---|---|---|---|---|---|---|
| Björk_6w | F | wolf | 2014 | Rolled up mattress | 24.939 | 7.938 | |
| Björk_10w | F | wolf | 2014 | Wheelbarrow | 30.704 | | 17.413 |
| Björk_14w | F | wolf | 2014 | Mirror | 67.582 | 3.071 | 14.622 |
| Björk_18w | F | wolf | 2014 | Stuffed wolverine | 14.082 | 1.027 | 5.11 |
| Björk_22w | F | wolf | 2014 | Mechanical dog | 13.722 | 5.399 | 3.343 |
| Björk_26w | F | wolf | 2014 | Moving sheet | 10.719 | 25.478 | 2.054 |
| Iggy_6w | M | wolf | 2014 | Rolled up mattress | 28.953 | 2.335 | 12.277 |
| Iggy_10w | M | wolf | 2014 | Wheelbarrow | 23.042 | 1.267 | 10.744 |
| Iggy_14w | M | wolf | 2014 | Mirror | 38.171 | 4.642 | 12.507 |
| Iggy_18w | M | wolf | 2014 | Stuffed wolverine | 22.75 | 2.778 | 71.645 |
| Iggy_22w | M | wolf | 2014 | Mechanical dog | 19.977 | 71.929 | 7.172 |
| Iggy_26w | M | wolf | 2014 | Moving sheet | 6.9 | 42.967 | 67.567 |
| Joni_6w | F | wolf | 2014 | Rolled up mattress | 42.269 | | 13.576 |
| Joni_10w | F | wolf | 2014 | Wheelbarrow | 113.923 | 31 | 66.565 |
| Joni_14w | F | wolf | 2014 | Mirror | 43.519 | 5.369 | 13.05 |
| Joni_18w | F | wolf | 2014 | Stuffed wolverine | 10.241 | 8.647 | 163.715 |
| Joni_22w | F | wolf | 2014 | Mechanical dog | 37.632 | 6.645 | 265.54 |
| Joni_26w | F | wolf | 2014 | Moving sheet | 1.295 | 208.822 | 25.069 |
| Lita_6w | F | wolf | 2014 | Rolled up mattress | 36.645 | 5.01 | |
| Lita_10w | F | wolf | 2014 | Wheelbarrow | 37.869 | 3.097 | 10.243 |
| Lita_14w | F | wolf | 2014 | Mirror | 21.78 | 8.702 | 5.139 |
| Lita_18w | F | wolf | 2014 | Stuffed wolverine | 11.534 | 4.133 | 45 |
| Lita_22w | F | wolf | 2014 | Mechanical dog | 20.482 | 3.698 | 14.848 |
| Lita_26w | F | wolf | 2014 | Moving sheet | 19.64 | 15.633 | 31.27 |
| Ozzy_6w | M | wolf | 2014 | Rolled up mattress | 42.991 | 4.608 | 4.118 |
| Ozzy_10w | M | wolf | 2014 | Wheelbarrow | 22.548 | 3.598 | |
| Ozzy_14w | M | wolf | 2014 | Mirror | 28.69 | 10.57 | |
| Ozzy_18w | M | wolf | 2014 | Stuffed wolverine | 7.977 | 2.298 | 42.982 |
| Ozzy_22w | M | wolf | 2014 | Mechanical dog | 2.039 | 17.907 | 191.145 |
| Ozzy_26w | M | wolf | 2014 | Moving sheet | 12.282 | 72.352 | 2.301 |
| Billie_6w | F | dog | 2015 | Rolled up mattress | 35.036 | 1.269 | 236.509 |
| Billie_10w | F | dog | 2015 | Wheelbarrow | 64.895 | 10.714 | |
| Billie_14w | F | dog | 2015 | Mirror | 228.046 | 6.404 | 104.38 |
| Billie_18w | F | dog | 2015 | Stuffed wolverine | 12.892 | 8.7 | 155.144 |
| Billie_22w | F | dog | 2015 | Mechanical dog | | 156.675 | |
| Billie_26w | F | dog | 2015 | Moving sheet | 1.004 | 287.973 | 7.706 |
| Ella_6w | F | dog | 2015 | Rolled up mattress | 43.391 | 12.308 | |
| Ella_10w | F | dog | 2015 | Wheelbarrow | 83.723 | 12.464 | |
| Ella_14w | F | dog | 2015 | Mirror | 277.401 | 2.037 | 57.007 |
| Ella_18w | F | dog | 2015 | Stuffed wolverine | 13.086 | 16.419 | 86.028 |
| Ella_22w | F | dog | 2015 | Mechanical dog | 13.302 | 7.408 | 1.772 |
| Ella_26w | F | dog | 2015 | Moving sheet | 2.031 | 21.754 | 15.08 |

*Table S2, continued*

| Individual | Sex | Species | Litter | Novel object | Invest_NO | Look_NO | Manip_NO |
|---|---|---|---|---|---|---|---|
| Muddy_6w | M | dog | 2015 | Rolled up mattress | 64.642 | 0.507 | 189.218 |
| Muddy_10w | M | dog | 2015 | Wheelbarrow | 71.701 | 30.783 | 3.071 |
| Muddy_14w | M | dog | 2015 | Mirror | 184.929 | 9.733 | 50.667 |
| Muddy_18w | M | dog | 2015 | Stuffed wolverine | 15.357 | 3.596 | 18.95 |
| Muddy_22w | M | dog | 2015 | Mechanical dog | 8.17 | 33.063 | |
| Muddy_26w | M | dog | 2015 | Moving sheet | 11.05 | 289.288 | 2.038 |
| Red_6w | M | dog | 2015 | Rolled up mattress | 56.028 | 6.414 | |
| Red_10w | M | dog | 2015 | Wheelbarrow | 38.345 | 7.431 | |
| Red_14w | M | dog | 2015 | Mirror | 104.427 | 1.267 | 15.108 |
| Red_18w | M | dog | 2015 | Stuffed wolverine | 8.182 | | 599.183 |
| Red_22w | M | dog | 2015 | Mechanical dog | 12.549 | 2.175 | 486.048 |
| Red_26w | M | dog | 2015 | Moving sheet | 1.794 | 12.984 | 9.943 |
| Simone_6w | F | dog | 2015 | Rolled up mattress | 114.982 | 0.501 | 100.587 |
| Simone_10w | F | dog | 2015 | Wheelbarrow | 86.153 | 1.533 | 27.381 |
| Simone_14w | F | dog | 2015 | Mirror | 135.977 | 7.116 | 71.895 |
| Simone_18w | F | dog | 2015 | Stuffed wolverine | 6.137 | | 37.77 |
| Simone_22w | F | dog | 2015 | Mechanical dog | 10.222 | 2.033 | |
| Simone_26w | F | dog | 2015 | Moving sheet | 3.568 | 13.074 | 3.331 |
| Skip_6w | M | dog | 2015 | Rolled up mattress | 95.184 | 1.03 | 44 |
| Skip_10w | M | dog | 2015 | Wheelbarrow | 98.82 | 37.07 | |
| Skip_14w | M | dog | 2015 | Mirror | 266.683 | 5.371 | 161.066 |
| Skip_18w | M | dog | 2015 | Stuffed wolverine | 35.395 | 12.328 | |
| Skip_22w | M | dog | 2015 | Mechanical dog | 13.784 | 5.591 | |
| Skip_26w | M | dog | 2015 | Moving sheet | 41.025 | 57.392 | 4.12 |
| Flea_6w | M | wolf | 2015 | Rolled up mattress | 86.793 | 14.762 | 31.947 |
| Flea_10w | M | wolf | 2015 | Wheelbarrow | 14.101 | 115.271 | |
| Flea_14w | M | wolf | 2015 | Mirror | 107.52 | 38.597 | 9.999 |
| Flea_18w | M | wolf | 2015 | Stuffed wolverine | 22.502 | 69.188 | 4.121 |
| Flea_22w | M | wolf | 2015 | Mechanical dog | 24.786 | 60.259 | |
| Flea_26w | M | wolf | 2015 | Moving sheet | 0.766 | 20.149 | 14.346 |
| Hendrix_6w | M | wolf | 2015 | Rolled up mattress | 10.238 | | 52.93 |
| Hendrix_10w | M | wolf | 2015 | Wheelbarrow | 51.255 | 4.338 | |
| Hendrix_14w | M | wolf | 2015 | Mirror | 75.441 | 13.077 | 36.64 |
| Hendrix_18w | M | wolf | 2015 | Stuffed wolverine | 15.321 | 1.798 | 12.602 |
| Hendrix_22w | M | wolf | 2015 | Mechanical dog | 11.24 | 4.5 | |
| Hendrix_26w | M | wolf | 2015 | Moving sheet | 14.572 | 13.285 | 36.652 |
| Elvis_6w | M | wolf | 2016 | Rolled up mattress | 70.845 | 22.34 | 21.968 |
| Elvis_10w | M | wolf | 2016 | Wheelbarrow | 6.903 | 74.034 | |
| Elvis_14w | M | wolf | 2016 | Mirror | 110.824 | 2.664 | 238.136 |
| Elvis_18w | M | wolf | 2016 | Stuffed wolverine | 48.436 | 8.52 | 7.411 |
| Elvis_22w | M | wolf | 2016 | Mechanical dog | 33.523 | 3.025 | 34.37 |
| Elvis_26w | M | wolf | 2016 | Moving sheet | | 18.881 | 14.103 |

*Table S2, continued*

| Individual | Sex | Species | Litter | Novel object | Invest_NO | Look_NO | Manip_NO |
|---|---|---|---|---|---|---|---|
| KD_6w | F | wolf | 2016 | Rolled up mattress | 73.791 | 29.971 | 85.252 |
| KD_10w | F | wolf | 2016 | Wheelbarrow | 65.794 | 69.81 | 14.842 |
| KD_14w | F | wolf | 2016 | Mirror | 193.343 | 12.792 | 127.961 |
| KD_18w | F | wolf | 2016 | Stuffed wolverine | 20.433 | 10.788 | 42.345 |
| KD_22w | F | wolf | 2016 | Mechanical dog | 34.494 | 38.522 | 7.291 |
| KD_26w | F | wolf | 2016 | Moving sheet | 15.489 | 73.536 | 6.357 |
| Lemmy_6w | M | wolf | 2016 | Rolled up mattress | 85.933 | 6.909 | 101.883 |
| Lemmy_10w | M | wolf | 2016 | Wheel barrel | | 95.109 | |
| Moby_6w | M | wolf | 2016 | Rolled up mattress | 46.788 | 31.431 | 107.534 |
| Moby_10w | M | wolf | 2016 | Wheelbarrow | 130.249 | 45.235 | |
| Moby_14w | M | wolf | 2016 | Mirror | 116.22 | 3.739 | 165.807 |
| Moby_18w | M | wolf | 2016 | Stuffed wolverine | | 240.913 | |
| Moby_22w | M | wolf | 2016 | Mechanical dog | 17.672 | 24.973 | |
| Moby_26w | M | wolf | 2016 | Moving sheet | | 31.539 | 13.52 |
| PJ_6w | F | wolf | 2016 | Rolled up mattress | 30.463 | | |
| PJ_10w | F | wolf | 2016 | Wheelbarrow | 33.27 | 14.137 | 22.518 |
| PJ_14w | F | wolf | 2016 | Mirror | 208.021 | | 212.876 |
| PJ_18w | F | wolf | 2016 | Stuffed wolverine | 7.582 | | 382.116 |
| PJ_22w | F | wolf | 2016 | Mechanical dog | 6.915 | 4.728 | 69.374 |
| PJ_26w | F | wolf | 2016 | Moving sheet | | 10.529 | 10.961 |
| Sting_6w | M | wolf | 2016 | Rolled up mattress | 99.796 | 8.515 | 55.313 |
| Sting_10w | M | wolf | 2016 | Wheelbarrow | 2.357 | 115.746 | |
| Sting_14w | M | wolf | 2016 | Mirror | 125.396 | 0.416 | 262.587 |
| Sting_18w | M | wolf | 2016 | Stuffed wolverine | 22.601 | 10.231 | 28.041 |
| Sting_22w | M | wolf | 2016 | Mechanical dog | 7.922 | 7.904 | 4.453 |
| Sting_26w | M | wolf | 2016 | Moving sheet | 0.315 | 13.486 | 9.644 |

**Table S3. Behavioural scores, non-novel object related.** Scores (seconds) for behaviours not related to the novel object for each of the six novel object tests.

| Individual | Sex | Species | Litter | Novel object | Active | Passive |
|---|---|---|---|---|---|---|
| Bowie_6w | M | dog | 2014 | Rolled up mattress | 142.296 | 389.466 |
| Bowie_10w | M | dog | 2014 | Wheelbarrow | 301.909 | 211.311 |
| Bowie_14w | M | dog | 2014 | Mirror | 280.560 | 269.010 |
| Bowie_18w | M | dog | 2014 | Stuffed wolverine | 251.085 | 282.194 |
| Bowie_22w | M | dog | 2014 | Mechanical dog | 366.421 | 169.383 |
| Bowie_26w | M | dog | 2014 | Moving sheet | 250.25 | 260.526 |
| Cash_6w | M | dog | 2014 | Rolled up mattress | 227.865 | 322.020 |
| Cash_10w | M | dog | 2014 | Wheelbarrow | 203.605 | 350.518 |
| Cash_14w | M | dog | 2014 | Mirror | 234.937 | 246.965 |
| Cash_18w | M | dog | 2014 | Stuffed wolverine | 275.517 | 282.37 |
| Cash_22w | M | dog | 2014 | Mechanical dog | 435.587 | 140.379 |
| Cash_26w | M | dog | 2014 | Moving sheet | 308.122 | 244.775 |
| Jagger_6w | M | dog | 2014 | Rolled up mattress | 224.256 | 310.453 |
| Jagger_10w | M | dog | 2014 | Wheelbarrow | 114.892 | 434.056 |
| Jagger_14w | M | dog | 2014 | Mirror | 80.983 | 266.718 |
| Jagger_18w | M | dog | 2014 | Stuffed wolverine | 343.138 | 232.931 |
| Jagger_22w | M | dog | 2014 | Mechanical dog | 165.902 | 273.522 |
| Jagger_26w | M | dog | 2014 | Moving sheet | 22.282 | 510.208 |
| Janis_6w | F | dog | 2014 | Rolled up mattress | 157.952 | 407.461 |
| Janis_10w | F | dog | 2014 | Wheelbarrow | 187.159 | 373.693 |
| Janis_14w | F | dog | 2014 | Mirror | 175.78 | 345.231 |
| Janis_18w | F | dog | 2014 | Stuffed wolverine | 154.308 | 278.274 |
| Janis_22w | F | dog | 2014 | Mechanical dog | 87.341 | 206.164 |
| Janis_26w | F | dog | 2014 | Moving sheet | 182.345 | 165.554 |
| Lennon_6w | M | dog | 2014 | Rolled up mattress | 216.505 | 337.187 |
| Lennon_10w | M | dog | 2014 | Wheelbarrow | 193.06 | 343.918 |
| Lennon_14w | M | dog | 2014 | Mirror | 232.568 | 289.471 |
| Lennon_18w | M | dog | 2014 | Stuffed wolverine | 228.667 | 318.318 |
| Lennon_22w | M | dog | 2014 | Mechanical dog | 252.897 | 50.351 |
| Lennon_26w | M | dog | 2014 | Moving sheet | 264.822 | 289.439 |
| Marley_6w | M | dog | 2014 | Rolled up mattress | 251.358 | 299.438 |
| Marley_10w | M | dog | 2014 | Wheelbarrow | 215.824 | 353.716 |
| Marley_14w | M | dog | 2014 | Mirror | 98.789 | 464.741 |
| Marley_18w | M | dog | 2014 | Stuffed wolverine | 32.33 | 243.667 |
| Marley_22w | M | dog | 2014 | Mechanical dog | 145.401 | 125.077 |
| Marley_26w | M | dog | 2014 | Moving sheet | 23.066 | 224.249 |

*Table S3, continued*

| Individual | Sex | Species | Litter | Novel object | Active | Passive |
|---|---|---|---|---|---|---|
| Björk_6w | F | wolf | 2014 | Rolled up mattress | 309.287 | 236.074 |
| Björk_10w | F | wolf | 2014 | Wheelbarrow | 330.614 | 215.092 |
| Björk_14w | F | wolf | 2014 | Mirror | 333.41 | 198.047 |
| Björk_18w | F | wolf | 2014 | Stuffed wolverine | 411.962 | 177.738 |
| Björk_22w | F | wolf | 2014 | Mechanical dog | 391.398 | 181.024 |
| Björk_26w | F | wolf | 2014 | Moving sheet | 346.281 | 130.024 |
| Iggy_6w | M | wolf | 2014 | Rolled up mattress | 315.331 | 235.283 |
| Iggy_10w | M | wolf | 2014 | Wheelbarrow | 361.502 | 209.256 |
| Iggy_14w | M | wolf | 2014 | Mirror | 371.669 | 185.999 |
| Iggy_18w | M | wolf | 2014 | Stuffed wolverine | 319.042 | 202.75 |
| Iggy_22w | M | wolf | 2014 | Mechanical dog | 299.115 | 155.695 |
| Iggy_26w | M | wolf | 2014 | Moving sheet | 210.961 | 79.845 |
| Joni_6w | F | wolf | 2014 | Rolled up mattress | 446.995 | 63.484 |
| Joni_10w | F | wolf | 2014 | Wheelbarrow | 320.604 | 64.579 |
| Joni_14w | F | wolf | 2014 | Mirror | 456.491 | 118.725 |
| Joni_18w | F | wolf | 2014 | Stuffed wolverine | 280.531 | 97.422 |
| Joni_22w | F | wolf | 2014 | Mechanical dog | 251.098 | 41.656 |
| Joni_26w | F | wolf | 2014 | Moving sheet | 225.494 | 57.851 |
| Lita_6w | F | wolf | 2014 | Rolled up mattress | 317.281 | 254.176 |
| Lita_10w | F | wolf | 2014 | Wheelbarrow | 367.552 | 133.813 |
| Lita_14w | F | wolf | 2014 | Mirror | 328.241 | 247.411 |
| Lita_18w | F | wolf | 2014 | Stuffed wolverine | 472.924 | 81.347 |
| Lita_22w | F | wolf | 2014 | Mechanical dog | 414.14 | 134.121 |
| Lita_26w | F | wolf | 2014 | Moving sheet | 113.098 | 54.285 |
| Ozzy_6w | M | wolf | 2014 | Rolled up mattress | 309.207 | 217.584 |
| Ozzy_10w | M | wolf | 2014 | Wheelbarrow | 354.186 | 231.048 |
| Ozzy_14w | M | wolf | 2014 | Mirror | 510.317 | 68.619 |
| Ozzy_18w | M | wolf | 2014 | Stuffed wolverine | 473.281 | 66.293 |
| Ozzy_22w | M | wolf | 2014 | Mechanical dog | 29.697 | 0 |
| Ozzy_26w | M | wolf | 2014 | Moving sheet | 328.035 | 202.478 |
| Billie_6w | F | dog | 2015 | Rolled up mattress | 175.355 | 165.004 |
| Billie_10w | F | dog | 2015 | Wheelbarrow | 200.115 | 322.793 |
| Billie_14w | F | dog | 2015 | Mirror | 129.36 | 133.17 |
| Billie_18w | F | dog | 2015 | Stuffed wolverine | 147.751 | 283.102 |
| Billie_22w | F | dog | 2015 | Mechanical dog | 272.825 | 149.434 |
| Billie_26w | F | dog | 2015 | Moving sheet | 87.424 | 230.613 |
| Ella_6w | F | dog | 2015 | Rolled up mattress | 226.538 | 323.645 |
| Ella_10w | F | dog | 2015 | Wheelbarrow | 334.23 | 166.245 |
| Ella_14w | F | dog | 2015 | Mirror | 137.554 | 74.593 |
| Ella_18w | F | dog | 2015 | Stuffed wolverine | 268.578 | 190.812 |
| Ella_22w | F | dog | 2015 | Mechanical dog | 437.413 | 120.119 |
| Ella_26w | F | dog | 2015 | Moving sheet | 252.097 | 306.013 |

*Table S3, continued*

| Individual | Sex | Species | Litter | Novel object | Active | Passive |
|---|---|---|---|---|---|---|
| Muddy_6w | M | dog | 2015 | Rolled up mattress | 183.985 | 172.007 |
| Muddy_10w | M | dog | 2015 | Wheelbarrow | 302.825 | 204.557 |
| Muddy_14w | M | dog | 2015 | Mirror | 271.787 | 86.725 |
| Muddy_18w | M | dog | 2015 | Stuffed wolverine | 304.048 | 268.572 |
| Muddy_22w | M | dog | 2015 | Mechanical dog | 347.387 | 229.627 |
| Muddy_26w | M | dog | 2015 | Moving sheet | 137.575 | 118.116 |
| Red_6w | M | dog | 2015 | Rolled up mattress | 132.019 | 408.392 |
| Red_10w | M | dog | 2015 | Wheelbarrow | 181.521 | 381.893 |
| Red_14w | M | dog | 2015 | Mirror | 226.518 | 248.825 |
| Red_18w | M | dog | 2015 | Stuffed wolverine | 0 | 0 |
| Red_22w | M | dog | 2015 | Mechanical dog | 27.054 | 88.615 |
| Red_26w | M | dog | 2015 | Moving sheet | 365.456 | 212.435 |
| Simone_6w | F | dog | 2015 | Rolled up mattress | 124.314 | 276.682 |
| Simone_10w | F | dog | 2015 | Wheelbarrow | 358.881 | 127.5 |
| Simone_14w | F | dog | 2015 | Mirror | 186.052 | 187.158 |
| Simone_18w | F | dog | 2015 | Stuffed wolverine | 387.862 | 189.129 |
| Simone_22w | F | dog | 2015 | Mechanical dog | 499.094 | 102.357 |
| Simone_26w | F | dog | 2015 | Moving sheet | 360.406 | 226.171 |
| Skip_6w | M | dog | 2015 | Rolled up mattress | 126.445 | 332.59 |
| Skip_10w | M | dog | 2015 | Wheelbarrow | 194.366 | 240.694 |
| Skip_14w | M | dog | 2015 | Mirror | 110.092 | 65.243 |
| Skip_18w | M | dog | 2015 | Stuffed wolverine | 316.883 | 247.807 |
| Skip_22w | M | dog | 2015 | Mechanical dog | 457.673 | 137.407 |
| Skip_26w | M | dog | 2015 | Moving sheet | 274.248 | 227.882 |
| Flea_6w | M | wolf | 2015 | Rolled up mattress | 312.501 | 132.463 |
| Flea_10w | M | wolf | 2015 | Wheelbarrow | 251.697 | 124.757 |
| Flea_14w | M | wolf | 2015 | Mirror | 285.102 | 138.143 |
| Flea_18w | M | wolf | 2015 | Stuffed wolverine | 345.721 | 116.481 |
| Flea_22w | M | wolf | 2015 | Mechanical dog | 377.67 | 114.948 |
| Flea_26w | M | wolf | 2015 | Moving sheet | 32.511 | 15.909 |
| Hendrix_6w | M | wolf | 2015 | Rolled up mattress | 138.618 | 390.782 |
| Hendrix_10w | M | wolf | 2015 | Wheelbarrow | 218.726 | 305.184 |
| Hendrix_14w | M | wolf | 2015 | Mirror | 244.397 | 233.58 |
| Hendrix_18w | M | wolf | 2015 | Stuffed wolverine | 330.623 | 259.577 |
| Hendrix_22w | M | wolf | 2015 | Mechanical dog | 473.868 | 178.163 |
| Hendrix_26w | M | wolf | 2015 | Moving sheet | 283.79 | 134.824 |
| Elvis_6w | M | wolf | 2016 | Rolled up mattress | 244.733 | 250.799 |
| Elvis_10w | M | wolf | 2016 | Wheelbarrow | 342.56 | 170.195 |
| Elvis_14w | M | wolf | 2016 | Mirror | 188.81 | 75.914 |
| Elvis_18w | M | wolf | 2016 | Stuffed wolverine | 370.331 | 181.448 |
| Elvis_22w | M | wolf | 2016 | Mechanical dog | 384.406 | 141.025 |
| Elvis_26w | M | wolf | 2016 | Moving sheet | 48.049 | 34.143 |

*Table S3, continued*

| Individual | Sex | Species | Litter | Novel object | Active | Passive |
|---|---|---|---|---|---|---|
| KD_6w | F | wolf | 2016 | Rolled up mattress | 289.904 | 118.009 |
| KD_10w | F | wolf | 2016 | Wheelbarrow | 289.217 | 115.959 |
| KD_14w | F | wolf | 2016 | Mirror | 159.677 | 133.547 |
| KD_18w | F | wolf | 2016 | Stuffed wolverine | 292.995 | 217.81 |
| KD_22w | F | wolf | 2016 | Mechanical dog | 361.189 | 158.362 |
| KD_26w | F | wolf | 2016 | Moving sheet | 307.914 | 54.177 |
| Lemmy_6w | M | wolf | 2016 | Rolled up mattress | 304.722 | 100.247 |
| Lemmy_10w | M | wolf | 2016 | Wheelbarrow | 300.52 | 72.63 |
| Moby_6w | M | wolf | 2016 | Rolled up mattress | 236.52 | 194.079 |
| Moby_10w | M | wolf | 2016 | Wheelbarrow | 294.773 | 63.035 |
| Moby_14w | M | wolf | 2016 | Mirror | 199.795 | 127.798 |
| Moby_18w | M | wolf | 2016 | Stuffed wolverine | 246.043 | 153.462 |
| Moby_22w | M | wolf | 2016 | Mechanical dog | 367.147 | 187.665 |
| Moby_26w | M | wolf | 2016 | Moving sheet | 31.096 | 24.753 |
| PJ_6w | F | wolf | 2016 | Rolled up mattress | 367.021 | 180.968 |
| PJ_10w | F | wolf | 2016 | Wheelbarrow | 399.582 | 110.937 |
| PJ_14w | F | wolf | 2016 | Mirror | 113.377 | 67.835 |
| PJ_18w | F | wolf | 2016 | Stuffed wolverine | 130.596 | 80.988 |
| PJ_22w | F | wolf | 2016 | Mechanical dog | 269.428 | 74.316 |
| PJ_26w | F | wolf | 2016 | Moving sheet | 259.093 | 183.891 |
| Sting_6w | M | wolf | 2016 | Rolled up mattress | 280.838 | 149.282 |
| Sting_10w | M | wolf | 2016 | Wheelbarrow | 361.584 | 95.797 |
| Sting_14w | M | wolf | 2016 | Mirror | 145.668 | 76.903 |
| Sting_18w | M | wolf | 2016 | Stuffed wolverine | 406.636 | 135.894 |
| Sting_22w | M | wolf | 2016 | Mechanical dog | 473.474 | 99.104 |
| Sting_26w | M | wolf | 2016 | Moving sheet | 109.991 | 53.29 |

**Table S4. Model selection.** Model selection table for each behaviour, listing the degrees of freedom (df), Akaike's Information Criterion (AIC), the difference in AIC with the best model (delta), and which model was selected.

| Variable | Model | df | AIC | delta | selected |
|---|---|---|---|---|---|
| **Latency, approach** | species * age | 7 | 297.48 | 0.00 | * |
| | species * age + sex | 8 | 298.18 | 0.70 | |
| | species + age | 6 | 300.84 | 3.35 | |
| | species + age + sex | 7 | 301.46 | 3.98 | |
| **Latency, contact** | species + age | 6 | 275.98 | 0.00 | |
| | species | 5 | 275.99 | 0.01 | * |
| | species * age | 7 | 277.52 | 1.54 | |
| | species + age + sex | 7 | 277.95 | 1.96 | |
| | species * age + sex | 8 | 279.49 | 3.51 | |
| **Looking at NO** | species * age | 8 | 503.60 | 0.00 | * |
| | species * age + sex | 9 | 504.43 | 0.83 | |
| | species + age | 7 | 505.94 | 2.35 | |
| | species + age + sex | 8 | 506.61 | 3.01 | |
| **Investigating NO** | species * age | 8 | 450.66 | 0.00 | * |
| | species * age + sex | 9 | 452.19 | 1.53 | |
| | species + age | 7 | 452.72 | 2.06 | |
| | species + age + sex | 8 | 454.10 | 3.44 | |
| **Manipulating NO** | species * age + sex | 9 | 599.78 | 0.00 | |
| | species * age | 8 | 599.91 | 0.14 | |
| | species + age + sex | 8 | 600.01 | 0.24 | |
| | species | 6 | 600.23 | 0.45 | * |
| | species + age | 7 | 600.41 | 0.63 | |
| **Active behavior** | species + age | 7 | 1755.21 | 0.00 | * |
| | species + age + sex | 8 | 1756.99 | 1.78 | |
| | species * age | 8 | 1757.20 | 2.00 | |
| | species | 6 | 1758.15 | 2.95 | |
| | species * age + sex | 9 | 1758.98 | 3.77 | |
| | species + sex | 7 | 1759.83 | 4.62 | |
| **Passive behavior** | species + age | 7 | 1698.84 | 0.00 | * |
| | species * age | 8 | 1699.36 | 0.52 | |
| | species + age + sex | 8 | 1700.51 | 1.67 | |
| | species * age + sex | 9 | 1700.96 | 2.12 | |
| | species | 6 | 1713.92 | 15.08 | |

**Table S5. Post hoc testing, contrast.** Pairwise species comparison at each age, based on models where age is a categorical factor. Post hoc testing was only performed for models were there was an overall interaction between species and age. Significant p-values are given in bold italic. P-values adjusted for multiple comparisons using Holm method are given.

| Behaviour | Contrast | Age | estimate | SE | df | t | p | adj. p |
|---|---|---|---|---|---|---|---|---|
| **Latency, approach** | *dog - wolf* | *6 weeks* | 0.096 | 0.291 | 17.558 | 0.330 | 0.745 | 1.000 |
| | *dog - wolf* | *10 weeks* | -0.407 | 0.291 | 17.558 | -1.399 | 0.179 | 0.896 |
| | *dog - wolf* | *14 weeks* | 0.004 | 0.295 | 18.666 | 0.014 | 0.989 | 1.000 |
| | *dog - wolf* | *18 weeks* | -0.384 | 0.295 | 18.666 | -1.301 | 0.209 | 0.896 |
| | *dog - wolf* | *22 weeks* | -0.246 | 0.295 | 18.666 | -0.834 | 0.415 | 1.000 |
| | *dog - wolf* | *26 weeks* | -0.923 | 0.295 | 18.666 | -3.131 | ***0.006*** | ***0.034*** |
| **Looking at NO** | *dog - wolf* | *6 weeks* | -0.540 | 0.545 | 10.783 | -0.991 | 0.343 | 1.000 |
| | *dog - wolf* | *10 weeks* | -0.519 | 0.544 | 10.769 | -0.953 | 0.361 | 1.000 |
| | *dog - wolf* | *14 weeks* | -0.162 | 0.552 | 11.57 | -0.292 | 0.775 | 1.000 |
| | *dog - wolf* | *18 weeks* | -0.182 | 0.552 | 11.533 | -0.329 | 0.748 | 1.000 |
| | *dog - wolf* | *22 weeks* | 0.612 | 0.558 | 11.934 | 1.097 | 0.294 | 1.000 |
| | *dog - wolf* | *26 weeks* | -0.113 | 0.644 | 19.888 | -0.176 | 0.862 | 1.000 |
| **Investigating NO** | *dog - wolf* | *6 weeks* | -0.140 | 0.411 | 25.481 | -0.342 | 0.735 | 1.000 |
| | *dog - wolf* | *10 weeks* | 0.741 | 0.410 | 25.454 | 1.805 | 0.083 | 0.415 |
| | *dog - wolf* | *14 weeks* | 0.341 | 0.417 | 27.225 | 0.818 | 0.421 | 1.000 |
| | *dog - wolf* | *18 weeks* | -0.222 | 0.417 | 27.116 | -0.533 | 0.599 | 1.000 |
| | *dog - wolf* | *22 weeks* | -1.196 | 0.422 | 28.029 | -2.831 | ***0.008*** | *0.051* |
| | *dog - wolf* | *26 weeks* | -0.300 | 0.504 | 45.615 | -0.596 | 0.554 | 1.000 |

**Table S6. Post hoc testing, slopes.** Slopes (trend) for each species for models where the interaction between species and age are significant. Lower and upper confidence intervals are given and significant confidence intervals are in bold italic.

| Behaviour | Species | Trend | SE | df | low.CI | up.CI |
|---|---|---|---|---|---|---|
| **Latency, approach** | *Dog* | -0.055 | 0.011 | 119.338 | ***-0.076*** | ***-0.033*** |
| | *Wolf* | -0.019 | 0.011 | 122.541 | -0.04 | 0.002 |
| **Looking at NO** | *Dog* | 0.121 | 0.021 | 118.168 | ***0.08*** | ***0.162*** |
| | *Wolf* | 0.057 | 0.024 | 122.963 | ***0.01*** | ***0.104*** |
| **Investigating NO** | *Dog* | -0.119 | 0.019 | 118.43 | ***-0.157*** | ***-0.082*** |
| | *Wolf* | -0.063 | 0.021 | 125.115 | ***-0.105*** | ***-0.021*** |

**Table S7. Model selection, latency to make contact (alternative analysis).** Model selection table for each behavior, listing the degrees of freedom (df), Akaike's Information Criterion (AIC), the difference in AIC with the best model (delta), and which model was selected.

| model | df | AIC | delta | Selected |
|---|---|---|---|---|
| *species + age* | 6 | 218.39 | 0 | |
| *species * age* | 7 | 219 | 0.61 | |
| *species + age + sex* | 7 | 219.54 | 1.15 | |
| *species* | 5 | 219.92 | 1.54 | * |
| *species * age + sex* | 8 | 220.17 | 1.78 | |

**Table S8. Model summary, latency to make contact (alternative analysis).** Results for the best fitted model of repeated measures, with dogs as the reference, on latency to make contact with the novel object, using lag time from the novel object was approached to the test ended as a measure latency for puppies that did not make contact with the novel object. Estimate, standard error, degrees of freedom, t-value and p-value are given. Significant p-values are marked in bold italic.

| Term | Estimate | Std. Error | df | t value | Pr(>\|t\|) |
|---|---|---|---|---|---|
| *(Intercept)* | 1.379 | 0.173 | 143.998 | 7.956 | ***0*** |
| *Species* | 0.169 | 0.243 | 143.999 | 0.692 | 0.49 |

**Table S9. Post hoc testing, latency to make contact (alternative analysis).** Pairwise species comparison at each age, based on models where age is a categorical factor. Significant p-values are given in bold italic. P-values adjusted (adj. p)for multiple comparisons using Holm method are given.

| Contrast | Age | Estimate | SE | df | t | p | adj. p |
|---|---|---|---|---|---|---|---|
| *dog - wolf* | 6 | -0.077 | 0.199 | 41.94 | -0.387 | 0.701 | 1 |
| *dog - wolf* | 10 | -0.156 | 0.199 | 41.94 | -0.781 | 0.439 | 1 |
| *dog - wolf* | 14 | 0.003 | 0.203 | 44.845 | 0.013 | 0.99 | 1 |
| *dog - wolf* | 18 | 0.118 | 0.203 | 44.845 | 0.584 | 0.562 | 1 |
| *dog - wolf* | 22 | 0.428 | 0.203 | 44.845 | 2.113 | ***0.04*** | 0.241 |
| *dog - wolf* | 26 | -0.069 | 0.203 | 44.845 | -0.342 | 0.734 | 1 |

**Table S10. Model selection, survival analyses of latency measures.** Model selection table for survival analyses on latency to approach and latency to contact the novel object, listing the degrees of freedom (df), Akaike's Information Criterion (AIC), the difference in AIC with the best model (delta), and which model was selected.

| Variable | model | df | AIC | delta | Selected |
|---|---|---|---|---|---|
| **Latency, approach** | species * age + sex | 12.09 | 1141.03 | 0 | |
| | species * age | 13.57 | 1141.99 | 0.97 | * |
| | species + age + sex | 9.52 | 1144.56 | 3.53 | |
| | species + age | 10.87 | 1145.33 | 4.31 | |
| | species + sex | 4.73 | 1154.46 | 13.43 | |
| | species | 6.18 | 1155.22 | 14.2 | |
| **Latency, contact** | species * age | 3 | 1116.12 | 0 | |
| | species + age | 2 | 1116.85 | 0.73 | * |
| | species * age + sex | 4 | 1118.04 | 1.92 | |
| | species + age + sex | 3 | 1118.8 | 2.68 | |
| | species | 1.01 | 1122.63 | 6.51 | |
| | species + sex | 2.01 | 1124.58 | 8.46 | |

**Table S11. Model summary, survival analyses of latency measures.** Results for the best fitted model of repeated measures, with dogs as the reference, on survival analyses on latency to approach and latency to make contact with the novel object. Estimate, standard error, z-value and p-value are given for each term. Significant p-values are marked in bold italic.

| *Behaviour* | *Term* | *Estimate* | *Std. error* | *z* | *p* |
|---|---|---|---|---|---|
| Latency, approach | *species* | -0.47 | 0.272 | -1.73 | 0.084 |
| | *age* | 0.08 | 0.021 | 3.77 | ***0*** |
| | *species:age* | -0.056 | 0.027 | -2.04 | ***0.041*** |
| Latency, contact | *species* | -0.211 | 0.173 | -1.22 | 0.22 |
| | *age* | -0.035 | 0.013 | -2.79 | ***0.005*** |

**Table S12. Model selection, excluding data for week 14.** Model selection table for each behaviour for models excluding week 14. Listed are the degrees of freedom (df), Akaike's Information Criterion (AIC), the difference in AIC with the best model (delta), and which model was selected.

| Behaviour | Model | df | AIC | delta | selected |
|---|---|---|---|---|---|
| **Latency, approach** | species * age | 7 | 255.86 | 0 | * |
| | species * age + sex | 8 | 256.06 | 0.2 | |
| | species + age + sex | 7 | 258.66 | 2.8 | |
| | species + age | 7 | 260.55 | 4.69 | |
| **Latency, contact** | species | 5 | 242.23 | 0 | * |
| | species + age | 6 | 243.2 | 0.97 | |
| | species * age | 7 | 244.91 | 2.68 | |
| | species + age + sex | 7 | 245.17 | 2.94 | |
| | species * age + sex | 8 | 246.88 | 4.65 | |
| **Look at NO** | species * age | 8 | 427.53 | 0 | |
| | species * age + sex | 9 | 428.35 | 0.82 | |
| | species + age | 7 | 429.03 | 1.5 | * |
| | species + age + sex | 8 | 429.7 | 2.17 | |
| | species | 6 | 455.2 | 27.66 | |
| **Investigating NO** | species * age | 8 | 356.14 | 0 | |
| | species + age | 7 | 357.26 | 1.12 | * |
| | species * age + sex | 9 | 357.79 | 1.65 | |
| | species + age + sex | 8 | 358.78 | 2.64 | |
| | species | 6 | 395.12 | 38.98 | |
| **Manipulating NO** | species | 6 | 499.45 | 0 | * |
| | species + age + sex | 8 | 499.7 | 0.26 | |
| | species * age + sex | 9 | 499.81 | 0.36 | |
| | species * age | 8 | 499.9 | 0.45 | |
| | species + age | 7 | 500 | 0.56 | |
| **Active behaviour** | species + age | 7 | 1460.14 | 0 | * |
| | species + age + sex | 8 | 1461.62 | 1.48 | |
| | species * age | 8 | 1462.13 | 1.99 | |
| | species | 6 | 1462.93 | 2.79 | |
| | species * age + sex | 9 | 1463.62 | 3.48 | |
| | species + sex | 7 | 1464.26 | 4.12 | |
| **Passive behaviour** | species + age | 7 | 1418.38 | 0 | * |
| | species * age | 8 | 1418.42 | 0.04 | |
| | species * age + sex | 9 | 1419.32 | 0.94 | |
| | species + age + sex | 8 | 1419.43 | 1.05 | |
| | species | 6 | 1434.62 | 16.24 | |

**Table S13. Model summary, excluding data for week 14.** Results for the best fitted model of repeated measures, with dogs as the reference, for each behaviour excluding data from week 14. Estimate, standard error, degrees of freedom, t-value and p-value are given. Significant p-values are marked in bold italic.

| *Behaviour* | *Term* | *Estimate* | *Std. error* | *df* | *t* | *p* |
|---|---|---|---|---|---|---|
| **Latency, approach** | *(Intercept)* | 0.876 | 0.147 | 2.251 | 5.965 | ***0.020*** |
| | *species* | 0.362 | 0.199 | 2.589 | 1.821 | 0.181 |
| | *age* | -0.055 | 0.011 | 94.650 | -4.982 | ***<0.0001*** |
| | *species:age* | 0.034 | 0.015 | 96.281 | 2.205 | ***0.030*** |
| **Latency, contact** | *(Intercept)* | 0.302 | 0.106 | 1.778 | 2.841 | 0.120 |
| | *species* | 0.295 | 0.146 | 2.116 | 2.014 | 0.175 |
| **Looking at NO** | *(Intercept)* | 1.314 | 0.836 | 54.034 | 1.573 | 0.122 |
| | *species* | 0.092 | 0.477 | 3.108 | 0.194 | 0.858 |
| | *age* | 0.091 | 0.016 | 96.768 | 5.757 | ***<0.0001*** |
| | *duration* | 0.126 | 0.074 | 99.855 | 1.695 | 0.093 |
| **Investigating NO** | *(Intercept)* | 1.036 | 0.633 | 98.765 | 1.637 | 0.105 |
| | *species* | 0.151 | 0.225 | 3.389 | 0.674 | 0.544 |
| | *age* | -0.089 | 0.013 | 96.804 | -6.836 | ***<0.0001*** |
| | *duration* | 0.175 | 0.060 | 106.353 | 2.912 | ***0.004*** |
| **Manipulating NO** | *(Intercept)* | 3.160 | 1.071 | 99.300 | 2.952 | ***0.004*** |
| | *species* | 0.463 | 0.402 | 2.641 | 1.150 | 0.344 |
| | *duration* | -0.129 | 0.101 | 117.723 | -1.272 | 0.206 |
| **Active behaviour** | *(Intercept)* | -168.095 | 56.145 | 113.464 | -2.994 | ***0.003*** |
| | *species* | 108.005 | 22.164 | 25.805 | 4.873 | ***<0.0001*** |
| | *age* | 2.476 | 1.128 | 97.994 | 2.195 | ***0.031*** |
| | *duration* | 38.816 | 5.281 | 102.913 | 7.350 | ***<0.0001*** |
| **Passive behaviour** | *(Intercept)* | 149.852 | 51.616 | 38.297 | 2.903 | ***0.006*** |
| | *species* | -95.001 | 32.964 | 2.928 | -2.882 | 0.065 |
| | *age* | -4.234 | 0.949 | 96.835 | -4.462 | ***<0.0001*** |
| | *duration* | 9.798 | 4.437 | 102.456 | 2.208 | ***0.029*** |

**Figure S1. Dog – wolf comparisons, excluding data for week 14.** Boxplots shows behavioural scores during a novel object test, comparing dogs and wolves across age. Overlaid are the fits and confidence intervals from the best model, selected by AIC. Boxes indicate the quartiles, and the whiskers reach maximally 1.5 times the interquartile range. Values beyond that are shown as points. An a log(x) scale) was used.

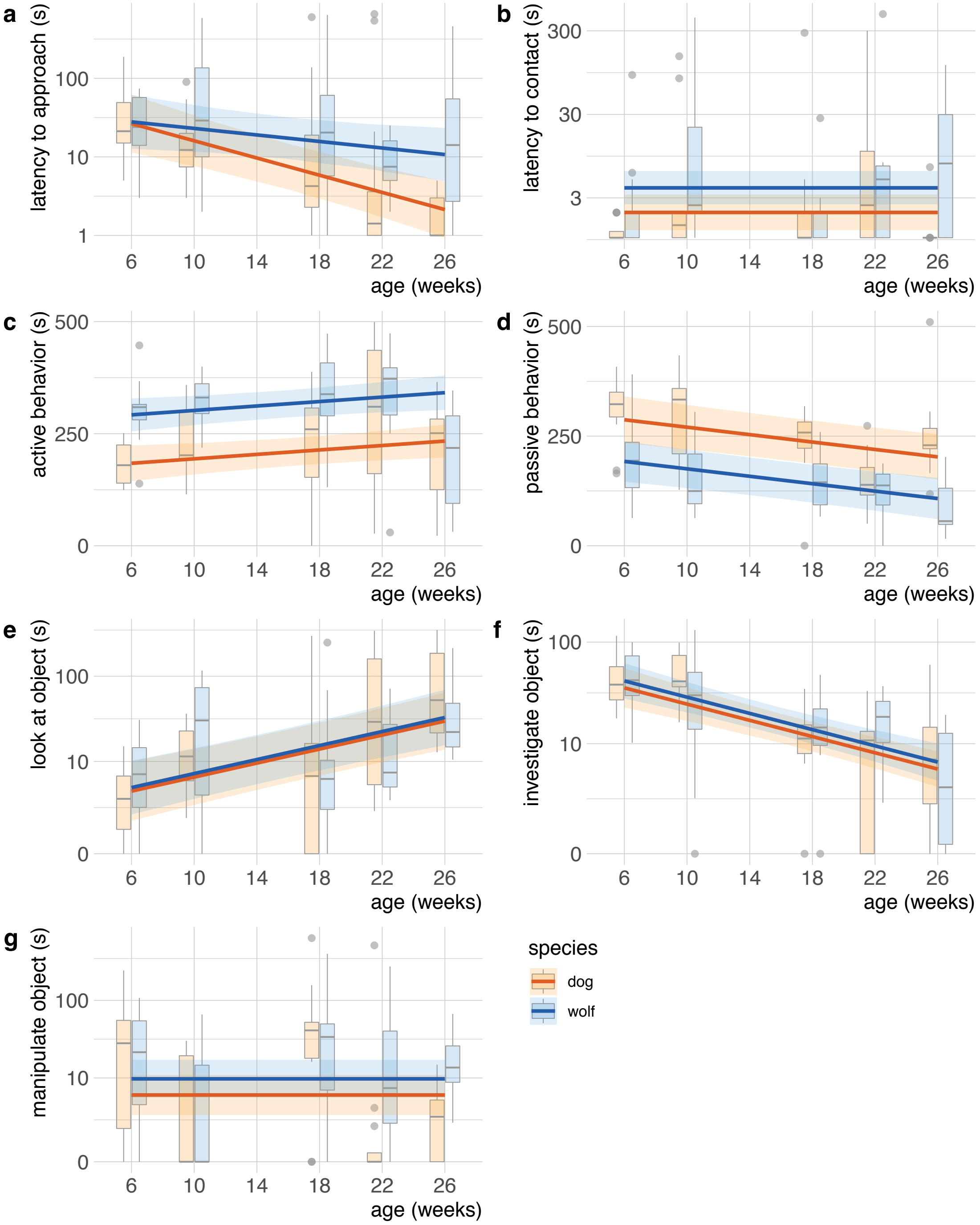

**Figure S2. Standardized regression coefficients, excluding data for week 14**. Standardized regression coefficients for the best model for each behaviour, excluding data from week 14, selected by AIC. Ranges indicate confidence intervals, computed using the likelihood profile. Missing estimates indicate that the term was not included in the best model.

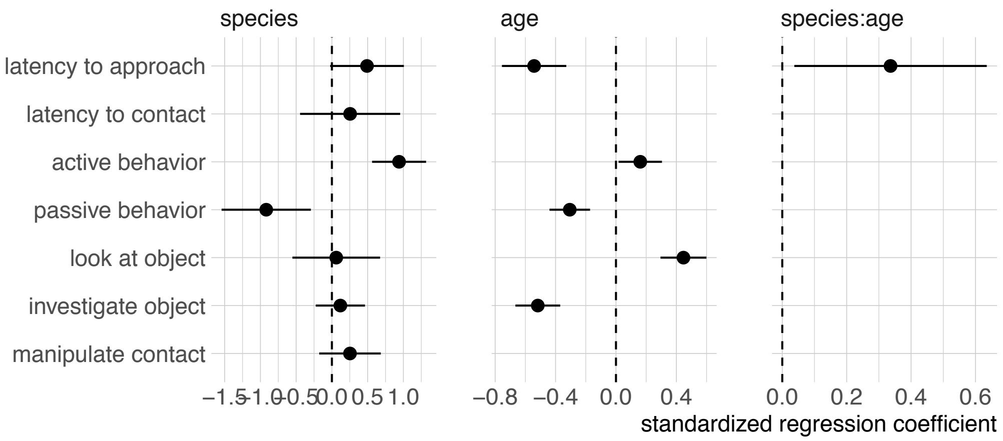
